# Translating the Post-Mortem Brain Multi-Omics Molecular Taxonomy of Alzheimer’s Dementia to Living Humans

**DOI:** 10.1101/2025.03.20.644323

**Authors:** Yasser Iturria-Medina, Victoria N. Poole, Andrea R. Zammit, Lei Yu, Shinya Tasaki, Joon Hwan Hong, Katia de Paiva Lopes, Caio Batalha, Abdur Raquib Ridwan, Ricardo A. Vialle, Lazaro Sanchez-Rodriguez, Maiya Rachel Geddes, Peter Abadir, Eric Ortlund, Philip De Jager, Vilas Menon, Michal Schnaider Beeri, Aron S. Buchman, Yishai Levin, David Morgenstern, Julie A. Schneider, Rima Kaddurah Daouk, Tony Wyss-Coray, Nicholas T. Seyfried, Konstantinos Arfanakis, Pedro Rosa-Neto, Yanling Wang, David A. Bennett

**Author notes:** Correspondence to: YIM and DAB.

## Abstract

Alzheimer’s disease (AD) dementia is characterized by significant molecular and phenotypic heterogeneity, which confounds its mechanistic understanding, diagnosis, and effective treatment. In this study, we harness the most comprehensive dataset of paired *ante-mortem* blood omics, clinical, psychological, and *post-mortem* brain multi-omics data and neuroimaging to extensively characterize and translate the molecular taxonomy of AD dementia to living individuals. First, utilizing a comprehensive integration of eight complementary molecular layers from brain multi-omics data (N = 1,189), we identified three distinct molecular AD dementia subtypes exhibiting strong associations with cognitive decline, sex, psychological traits, brain morphology, and characterized by specific cellular and molecular drivers involving immune, vascular, and oligodendrocyte precursor cells. Next, in a significant translational effort, we developed predictive models to convert these advanced brain-derived molecular profiles (AD dementia pseudotimes and subtypes) into blood-, MRI- and psychological traits-based markers. The translation results underscore both the promise of these models and the opportunities for further enhancement. Our findings enhance the understanding of AD heterogeneity, underscore the value of multi-scale molecular approaches for elucidating causal mechanisms, and lay the groundwork for the development of novel therapies in living persons that target multi-level brain molecular subtypes of AD dementia.

## INTRODUCTION

Recent advances in high-throughput tools for genome-wide multi-level omics analyses have facilitated the application of innovative molecular approaches to further characterize AD progression and heterogeneity (*1–6*). Independent studies (*3, 5–9*) have revealed distinct AD subgroups, suggesting subtype-specific molecular dysregulations associated with the progressive loss of cognition that eventually leads to AD dementia. Considering that the scope of molecular changes in AD extends beyond individual omic types, we recently proposed advanced machine-learning (ML) to unify four different molecular omics from *post-mortem* brain samples (*10*). This allowed us to detect distinct, molecularly-differentiable and statistically stable pseudotime subtrajectories (subtypes) from no cognitive impairment (NCI) to AD dementia, and to quantify person-specific progression along these subtrajectories (*10*). These subtypes represent three different molecular paths from no cognitive impairment (NCI) to AD dementia. In agreement with other AD molecular classifications (*3, 6, 7*), our results confirmed that the diversity of AD dementia cannot be fully accounted for by a single molecular omic, neuropathological patterns, clinical manifestations, or demographic variations. This highlighted the importance of considering complementary disease processes across different molecular levels. Hence, enhancement of such multi-scale approaches is critical to robustly uncover distinctive AD dementia pathways and meaningfully incorporate the wide array of molecular layers now being generated from *post-mortem* human brain samples.

Our prior study had some limitations. Most importantly, the deepest data was the bulk RNAseq followed by the DNA 5C methylation, with significantly fewer metabolomic features followed by even fewer targeted proteomics (17). Here, we leverage the deepest set of brain multi-omics data integrating eight complementary molecular layers in a relatively large number (up to N∼1,200). This includes DNA methylation, single-nucleus and bulk RNA sequencing, tandem mass tag (TMT) proteomic, glycated proteomic, glycosylated proteomic, metabolomic and lipidomic data from *post-mortem* brains, unified via advanced ML. In addition to updating our AD dementia pseudotime and molecular subtype results, we perform robust causal analysis to identify the cell-type specific genetic drivers of each distinctive molecular AD dementia subtype, clarifying specific disease mechanisms and identifying potential tailored therapeutic targets.

Next, we address one of the most challenging tasks in the field. Namely, translating the brain molecular omic taxonomy derived from tissue samples from decedents to living older adults. Here, we do not take an *in silico* approach. Rather, we leverage data collected *ante-mortem* from the same humans on whom we generated the brain molecular taxonomy. These data includes blood omics, AD blood biomarkers, routine blood tests, psychological, and neuroimaging data (in this case *ex vivo* as a proxy for *in vivo* MRI). Notably, the blood omics data incorporates cell-free DNA, cytokines, monocytes transcriptomic, proteomic, and metabolomic levels. From this multi-level data we begin to develop predictors of brain molecular AD dementia pseudotimes and subtypes in living people. Overall, our results chart a clear pathway to create disease predictors of deeply characterized brain molecular AD dementia pseudotime and subtypes in living persons. Thus, for the first time to our knowledge, we lay the groundwork for developing therapeutics that target AD dementia molecular trajectories generated from brain omics and translated to life.

## RESULTS

### Data origin and unifying analytical approach

We obtained paired multi-omics molecular data from *post-mortem* brains (N=1,189) and *in-vivo* blood (N=1,108) from the same ROSMAP participants with a wide range of cognition covering the continuum from no cognitive impairment to AD dementia (see Figs. 1A and Table S1; *Materials and Methods*, *Data*). Brain-derived data included DNAm (originally >65,000 CpG islands), snRNA (7 cell-types, >43,000 transcripts), bulk RNA-seq (∼17,300 genes), proteins (8,356 proteins), glycopeptiforms (11,012 total mapping to 1,326 protein groups), metabolites (735) and lipids (343) concentrations, along with structural MRI (*11*) and extensive neuropathological evaluations (*12–15*). Blood data included cell-free DNA (6 markers), monocytes RNA-seq (>52,000 transcripts), somalogic proteins (7,289), p-tau181 and p-tau217, cytokines (4), and metabolites (292). Clinical diagnoses were no cognitive impairment (NCI), mild cognitive impairment (MCI) or AD dementia proximate to death (*16–18*) (*Materials and Methods*, *Data;* Table S1). Five psychological traits were used, including depressive symptomatology, neuroticism, self-perceived loneliness, conscientiousness, and purpose in life (*19–24*).

**Figure 1.**
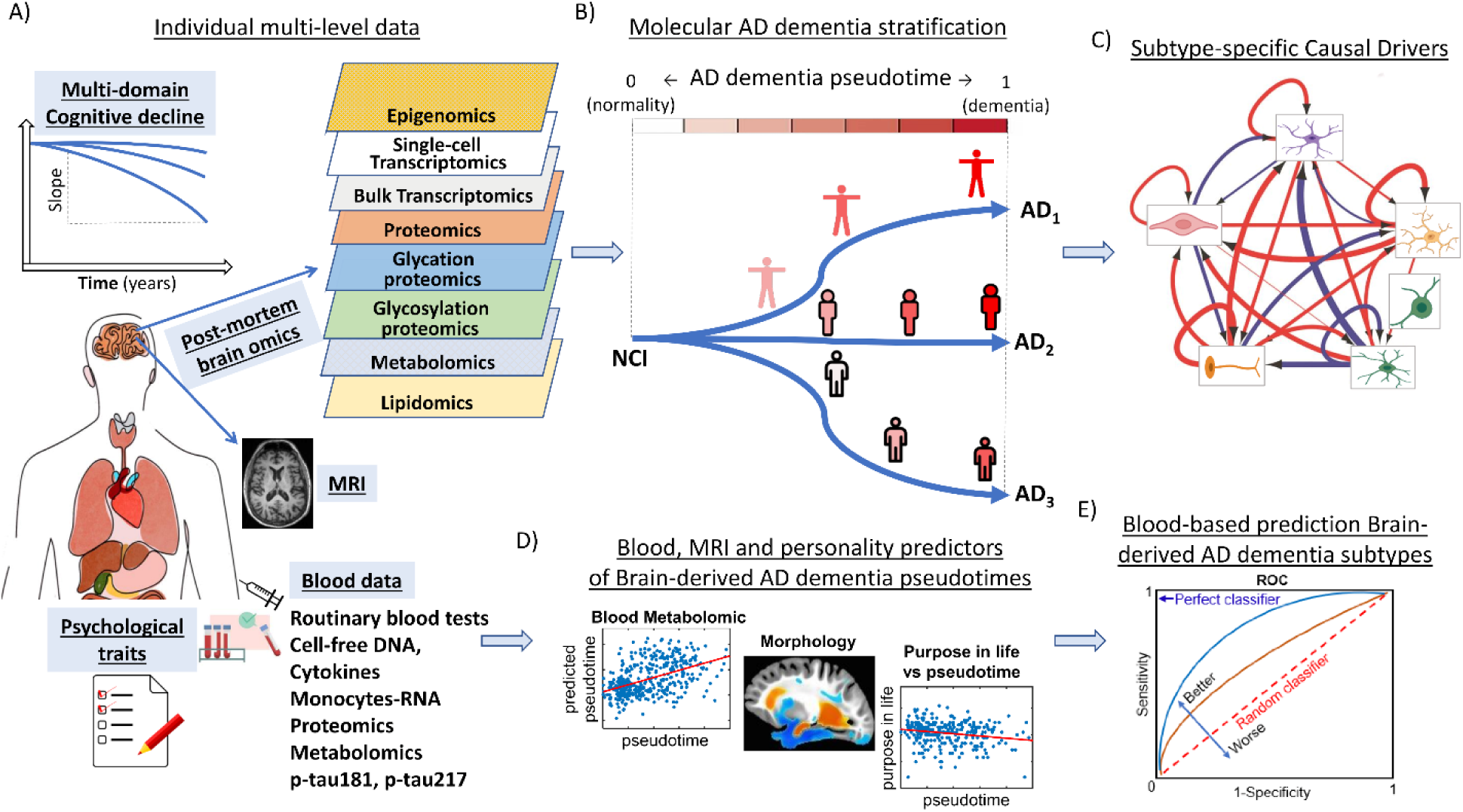
Schematic approach for identifying, characterizing and translating the brain multi-omics molecular AD dementia taxonomy. A) Multi-modal *ante-mortem* data including *in-vivo* blood, longitudinal multi-domain cognition, survey, and neuroimaging data, paired with *post-mortem* brain data, from the same ROSMAP participants. B) A contrastive machine-learning algorithm was used to integrate the brain’s multi-omics molecular data, obtaining a multidimensional disease space where subjects define biologically-distinctive subtrajectories from no cognitive impairment (NCI) to AD dementia. We first obtained for each participant an AD dementia pseudotime score and subsequently an AD dementia subtype. C) Subsequently, each AD dementia subtype was further characterized by identifying specific cell-type causal drivers and multi-omics features, providing insights into distinct disease mechanisms. D-E) Lastly, we tested whether brain-derived pseudotimes and AD dementia subtypes can be predicted by multi-modal data obtained from the same participants prior to death.

We proceeded to identify and characterize the brain multi-omics molecular taxonomy of AD dementia. To do so, we unified, reordered, and stratified the eight brain omic molecular data layers (Figs. 1A,B) using a recently proposed ML algorithm (*10*) for detection of disease-associated multimodal data patterns. The data were anchored in NCI, the preferred state, relative to AD dementia, the disease state, and MCI allowed to passively assort (*Materials and Methods, Multimodal contrastive Trajectories Inference [mcTI] definition*; Fig. 1A). All molecular markers were first adjusted for potentially confounding covariates (e.g., age, sex, education, *post-mortem* interval). Similar to our previous work (*10*), which was limited to four brain molecular omic layers, here we assumed that the position of each subject in the aggregated molecular space would predict individual severity of AD dementia and placement in distinctive disease brain molecular pseudotime subtypes. A molecular AD dementia pseudotime representing the biological “distance” or “progression” from NCI to AD dementia was calculated for each subject, ranging from 0 (indicating NCI) to 1 (AD dementia). Next, each participant (except NCI individuals) was assigned to a distinct AD dementia subtype based on the highest likelihood of aligning with characteristic disease patterns in the multi-scale molecular space (see Fig. 1B). Each AD dementia subtype represents a unique brain multi-omics molecular pathway from normality to AD dementia.

We next characterized each brain AD dementia subtype by identifying its cell-type specific causal drivers by inferring direct regulatory networks from subtype-specific snRNA data (Fig. 1C). Finally, we explore the ability of *in-vivo* data including blood omics, psychological traits, and neuroimaging (ex vivo as a proxy for in vivo) to translate the *post-mortem* brain multi-omics AD dementia subtypes (see schematic Figs. 1D, E) to data generated from the same humans prior to death.

### Characterization of AD dementia brain multi-omics molecular pseudotime

First, we confirm that the *post-mortem* brain multi-omics molecular AD dementia pseudotime predicts cognitive performance and decline. We observed strong negative associations between molecular AD dementia pseudotime and multiple cognitive domains at baseline and their rates of change over time leveraging the cognitive function data obtained over multiple years prior to death. (Fig. 2A; all P<0.05, FWE-corrected, based on correlation tests with permutations after adjustment by age, sex, and educational level). Strongest associations corresponded to global cognition and episodic memory, and temporal decline in global cognition, with higher pseudotime values representing faster cognitive decline, as expected.

**Figure 2.**
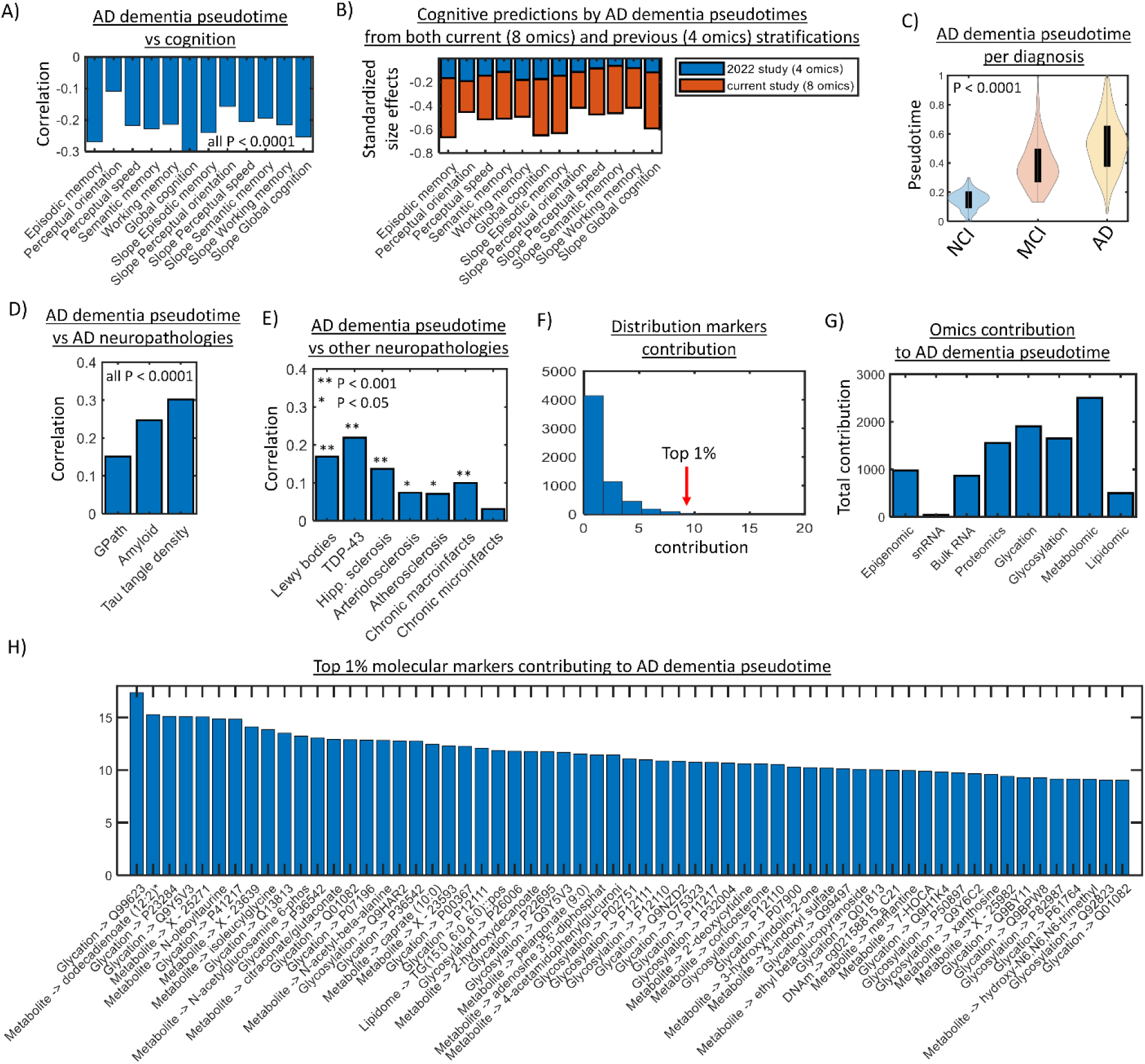
Characterization of multi-omics molecular AD dementia pseudotimes. A) AD dementia pseudotime correlation values with multiple cognitive domains and their rates of change over time (all P<0.05, FWE-corrected, based on correlation tests with permutations after adjustment by age, sex, and educational level). Notice the strong associations with episodic memory, global cognition, and their temporal declines. B) Standardized size effects for AD dementia pseudotimes, from the current study and our previous four-omics version from 2022 (*10*), when concurrently explaining cognitive performance and decline. For each cognitive variable, we conducted linear regression models incorporating both sets of AD dementia pseudotime as predictors along with age, sex, and educational level as covariables. C) AD dementia pseudotime distributions across clinical diagnoses (NCI, MCI, AD; p-value FWE-corrected, based on Ancova tests with age, sex, and educational level as covariables). D-E) AD dementia pseudotime correlation with AD and non-AD specific neuropathologies, respectively (FWE-corrected, based on correlation tests with permutations after adjustment by age, sex, and educational level). F) Distribution of all multi-omics markers’s contributions to AD dementia pseudotimes. G) Omic-specific contributions. H) Top 1% contributing omic markers.

To evaluate the added value of the additional four omic layers relative to the initial four-omics version generated by our team in 2022 (*10*), we assessed its additional predictive power concerning baseline cognitive performance and rates of temporal decline. Specifically, we conducted linear regression models for each cognitive variable, incorporating both AD dementia pseudotime sets (i.e., from our current and 2022 studies), along with age, sex, and educational level as covariates. Our analysis revealed a significantly stronger predictive performance for the updated AD dementia pseudotimes, evidenced by standardized beta coefficients that were approximately 2 to 4 times larger for each cognitive variable (see Fig. 2B; all P<0.001 from Ratio tests).

Based on ANCOVA tests with age, sex, and educational level as covariables, we also confirmed that the updated AD dementia pseudotime values were significantly associated with the individual clinical diagnosis (Fig. 2C; *P<0.0001*, FWE-corrected). Furthermore, the molecular AD dementia pseudotimes significantly correlated with landmark AD neuropathologies, including global burden of AD pathology, average amyloid-β load and PHF tau tangle density (Fig. 2D; P<0.0001, FWE-corrected, covariables adjusted). They also significantly correlated with the severity of non-AD neuropathologies, including Lewy bodies, TDP-43, hippocampal sclerosis, arteriolosclerosis, atherosclerosis, chronic macroinfarcts (Fig. 2E; all P<0.05, FWE-corrected, covariables adjusted), and did not correlate significantly with chronic microinfarcts. This is expected as AD dementia frequently co-occurs with other pathologies (*14, 15, 25, 26*).

Next, we quantified the contribution of all the brain molecular markers and omic types on the resulting AD dementia pseudotimes. The mcTI algorithm allowed us to analyze the loadings (or model weights) from the internal dimensionality reduction across data types and the subsequent principal components fusion, providing a direct quantification of the relevance of each specific omic marker and type on pseudotimes estimation (*Materials and Methods*, subsection *Assessing markers contributions on pseudotime*). Across markers, we observed a pronounced decay in contribution to the AD dementia pseudotimes, with about 4,000 markers (out of 6,085 total) relatively close to zero values (Fig. 2F). The pseudotime was predominantly informed by metabolomic, glycation, glycosylation and proteomic layers (Fig. 2G), with snRNA data exerting comparatively lesser impact. The top 1% contributing features (Fig. 2H) abounded in glycation, metabolomic and glycosylation markers. The identification of glycation markers as strong pseudotime contributors is noteworthy, as they—such as advanced glycation end products (AGEs)—can accumulate in neural tissues and are associated with neurodegeneration. For instance, the presence of AGEs in amyloid-β plaques and neurofibrillary tangles can exacerbate oxidative stress and neuronal death (*27, 28*). Metabolites like dodecadienoate and N-acetylglucosamine 6-phosphate also emerged as critical contributors. Dodecadienoate, a polyunsaturated fatty acid, may influence inflammatory pathways, while N-acetylglucosamine 6-phosphate is involved in cellular metabolism and signaling, potentially affecting neuronal health and function (*29, 30*). Other identified metabolites, such as isoleucylglycine and adenosine 3’,5’-diphosphate, could reflect altered energy metabolism and neurotransmitter dynamics in the context of cognitive decline (*30, 31*). The observed interplay between glycation, glycosylation and metabolite changes underscores the complexity of AD pathology. Dysregulation in glycation and glycosylation products not only disrupt normal cellular functions but also interacts with metabolic pathways that are crucial for maintaining neuronal integrity.

These findings support the idea that distinct biological data play varying roles in informing about the progression of AD dementia (*25, 32–34*). They also suggest that biological units with relatively lower-resolution (e.g., metabolites, glycoproteins) may be more informative for tracking AD dementia progression, given their stronger associations with clinical phenotypes. However, it is important to note that causality was not implied in this type of analysis; thus, the results should be interpreted in the context of quantifying disease progression rather than elucidating disease mechanisms. In this respect, data from high-resolution biological scales (e.g., snRNA) may be particularly informative, as studied below.

### Characterization of AD dementia multi-omics brain molecular subtypes

Our ML algorithm also assigned each individual to one of three distinct AD dementia molecular subtypes (Figs. 3A, B) by detecting unique multimodal patterns within the molecular landscape. In this context, AD dementia subtypes represent distinct molecular pathways from cognitive normality to AD dementia. To identify the optimum number of subtypes (from a minimum of 1 up to a maximum of 5), we used a majority rule across the Calinski–Harabasz, Davies-Bouldin, Gap, and Silhouette criteria (*35, 36*). The stability and significance of each subtype configuration were tested via a permutation procedure (*3*), evaluating the rate at which sample pairs group together into the same subtypes upon repeated clustering on random subsets.

**Figure 3.**
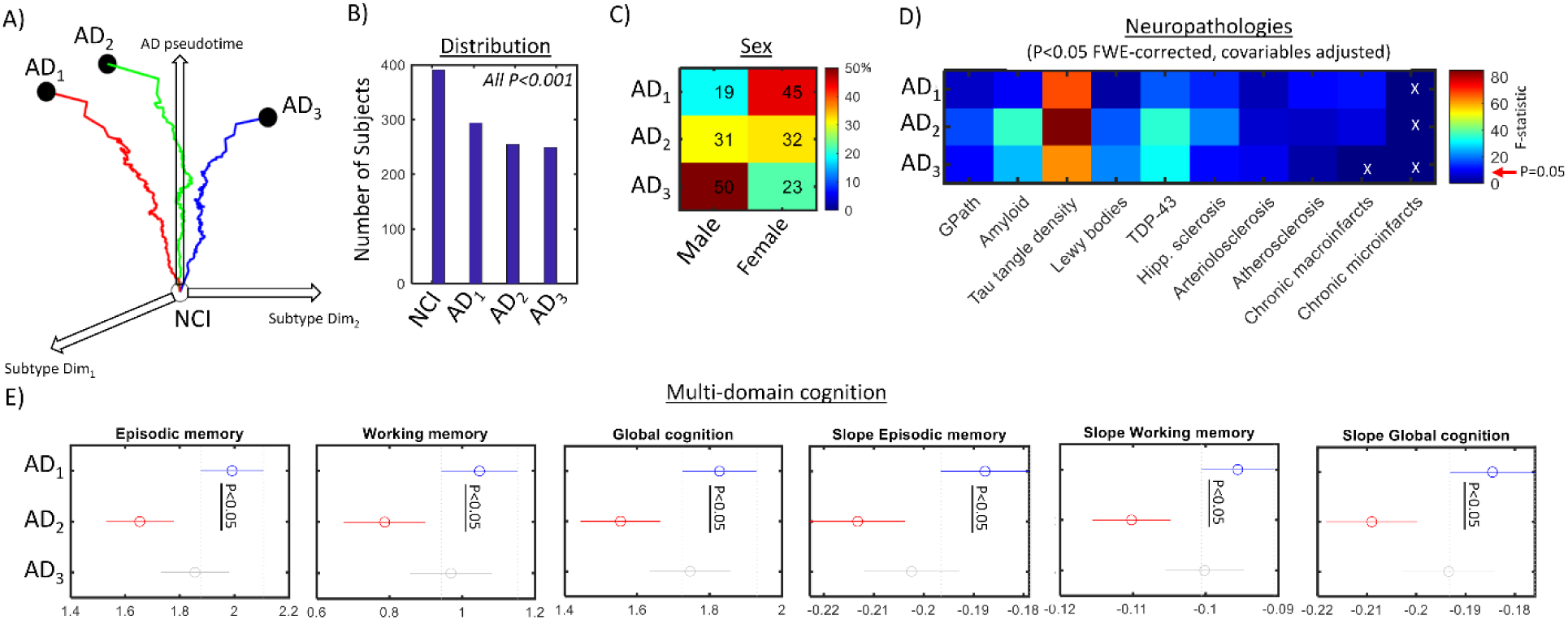
Distinctive multi-level AD dementia subtypes identified with *post-mortem* brain data. A) Three-dimensional visualization of the identified AD dementia subtypes’ subtrajectories from cognitive normality to AD dementia. X and Y axes correspond to the first two dimensions on the reduced multi-omics molecular space, and Z axis to the AD dementia pseudotime. B) Number of participants per AD subtype and background subpopulation (NCI), and corresponding significance obtained with randomization testing (all P<0.001, FWE-corrected). C) Subtype-specific sex proportions, normalizing across all the participants in the cohort. D) Subtype-specific statistical differences in severity of AD-related and non-AD neuropathologies, resulting from ANCOVA tests with subtype as grouping variable (color scale corresponds to F-values, FWE-corrected via permutation tests). E) Across subtypes comparison tests in cognitive performance. Tukey’s honest significant difference criterion was used (with P<0.05 implying significance differences in marginal means; *Materials and Methods*, *Additional Statistical Analysis*).

The three AD dementia molecular subtypes AD_1_, AD_2_ and AD_3_ had similar numbers of participants, i.e., 294, 255 and 249 (Fig. 3B). AD_2_ included a balanced distribution of female and male participants, comprising 31% and 32% of the total numbers in the cohort, respectively (see Fig. 3C). By contrast, AD_1_ was predominantly female, accounting for 45% of all females in the cohort but only 18% of the males, and AD_3_ had the opposite ’sex-dominance’ pattern, including 23% of the females and 50% of the males. As noted above, prior to the mcTI analysis, all molecular data were linearly adjusted for sex reducing the likelihood that the sex differences were the result of confounding. Thus, there appears to be two relatively sex-specific molecular pathways to AD dementia in addition to one unrelated to sex.

When compared with the reference subpopulation (NCI), the three subtypes exhibited substantial alterations in AD-related neuropathologies as expected, including greater overall AD pathologic burden, average amyloid-β load and PHFtau tangle density (Fig. 3D; all P<0.01, FWE-corrected, covariables adjusted). They also presented significant non-AD neuropathologies, such as Lewy bodies, TDP-43, hippocampal sclerosis, arteriolosclerosis, atherosclerosis, vascular infarcts (Fig. 3D; all P<0.05, FWE-corrected, covariables adjusted), except for AD_3_ in severity of vascular infarcts. The three subtypes did not show significant differences in the severity of chronic microinfarcts compared to NCI. While the three subtypes exhibited comparable neuropathological changes relative to NCI, AD_1_ stood out with a significantly lower presence of Lewy bodies compared to both AD_2_ and AD_3_, as well as significantly higher occurrence of vascular infarcts than AD_3_ (all P<0.05, Tukey’s honest significant difference test).

In addition, AD_2_ exhibited lower cognitive performance than AD_1_ in global cognition, specifically episodic memory and working memory, and their change over time (Fig. 3E; all P<0.05, Tukey’s honest significant difference test). The subtypes were not statistically different in perceptual orientation, perceptual speed, semantic memory, or their corresponding rates of change in time.

We proceeded to identify molecular alterations distinctively contributing to each AD dementia subtype. For this, we considered NCI as reference group to detect omic features significantly altered for each subtype (P<0.05, FDR-corrected) while statistically preserved in the other two. This resulted in subtype-specific molecular signatures (*Materials and Methods, Identification of Subtypes Signatures*). AD_1_’s signature was characterized by 71 uniquely differentiated omic markers, AD_2_ by 1035, and AD_3_ by 342. Figure 4A presents the top 30 distinctive contributors for each subtype (for a complete list per subtype, see *Files S1-S3*).

**Figure 4.**
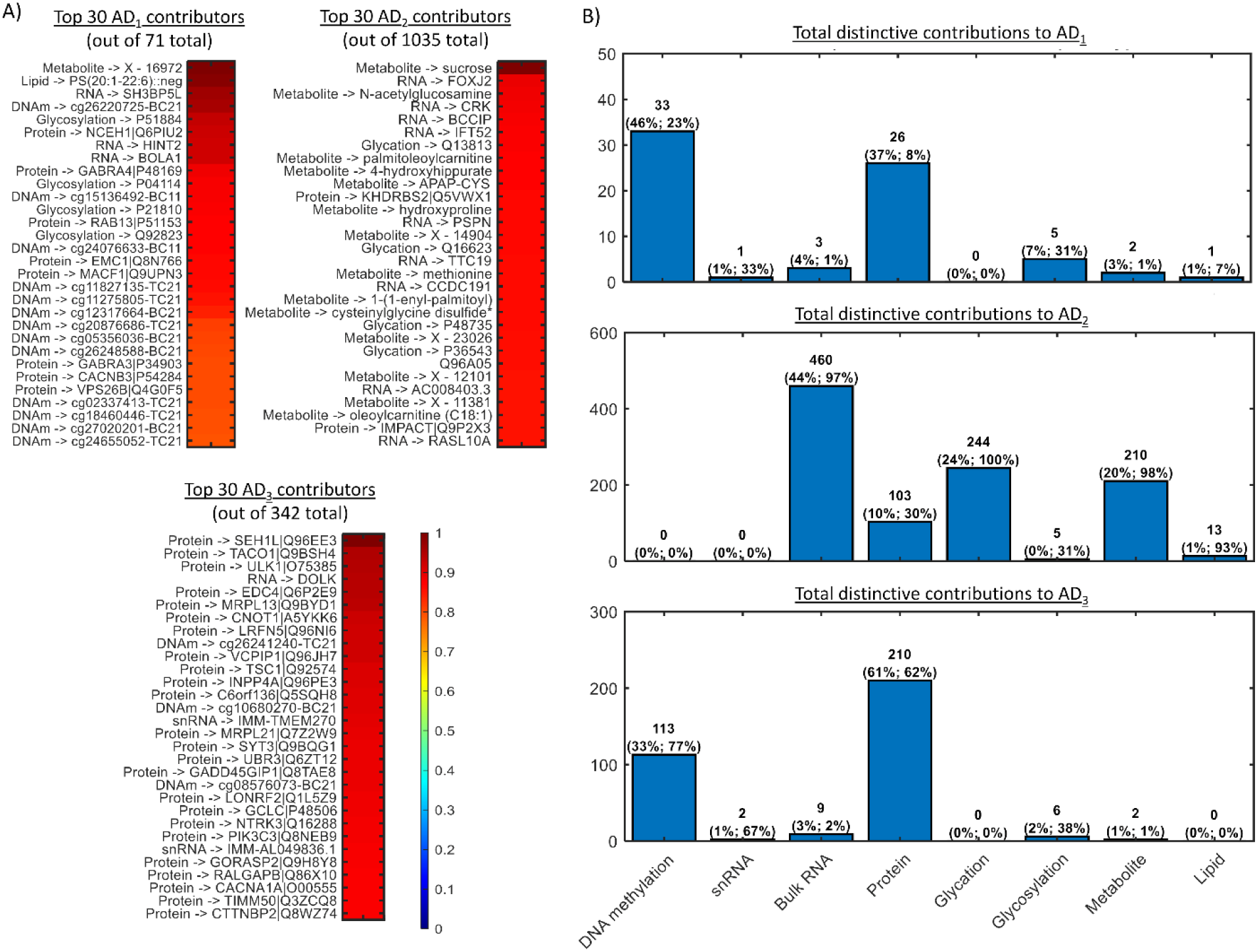
Subtypes-specific multi-level molecular alterations. A) Top 30 molecular features distinctively altered for each subtype while preserved in the other subtypes (for complete lists, see *Files S1-S3*). Contribution values per subtype are normalized by the maximum. B) The raw values displayed above the bars represent the unique counts of distinct contributors for each subtype across specific omics. The first reported percentage for omic type (i) in a given subtype (j) indicates the number of markers from omic type (i) that are uniquely altered for that subtype, normalized by the total number of uniquely altered markers for subtype (j) across all omics. The second percentage for omic type (i) in a subtype (j) reflects subtype-specific percentages of distinctive alterations across all subtypes. Specifically, it corresponds to the number of markers from omic type (i) that are uniquely altered for subtype (j), normalized by the total number of markers of omic type (i) uniquely altered across all subtypes.

Figure 4B presents the modality-specific number of distinct molecular contributors for each subtype. We performed two complementary analyses to further clarify these results. First, for each subtype, omic-specific percentages of distinctive alterations were calculated, i.e., normalizing within each subtype by its total number of distinctively altered omic features (see first percentage reported above each bar on Fig. 4B). We observed that AD_1_’s distinctive alterations were predominantly at the epigenomic (46%) and proteomic (37%) levels. AD_2_ was mostly defined by distinctive bulk-RNA (44%), glycated (24%) and metabolomic (20%) alterations, distributed across a total of 1,035 distinctive contributors. AD_3_ was predominantly differentiated at the proteomic (61%) and epigenomic (33%) scales.

Second, for each omic, subtype -specific percentages of distinctive alterations were calculated, i.e., normalizing within each omic type by the total number of distinctively altered features across subtypes (see second percentage reported above each bar on Fig. 4B). We observed that, across all subtypes, AD_2_ covered most of the unique alterations detected for the glycation (100%), metabolomic (98%), bulk RNA (97%) and lipidomic (93%) data layers. AD_3_ was primarily characterized by unique epigenomic (77%), snRNA (67%) and proteomic (62%) alterations. In turn, AD_1_ was associated with considerably fewer unique alterations across subtypes, been distinctively defined by snRNA (33%), epigenetic (31%) and glycosylation (23%) differences.

Overall, our findings indicate a relatively low degree of overlap in types of molecular alterations among the three AD dementia subtypes. Moreover, the results (Figs. 4A, B) suggest that AD_2_ may represent a more biologically aggressive variant of the disease at multiple molecular levels (with approximately 14 and 3 times more significantly altered molecular markers than AD_1_ and AD_3_, respectively), and characterized by significantly lower cognitive function (as shown in Figs. 4E).

We next quantified the total contribution of the molecular data types on the resulting AD dementia molecular subtypes. To achieve this, we iteratively repeated the subtrajectories-identification process after systematically excluding each individual data layer. This method allowed us to quantify the deviations from the original AD subtypes that occurred when each layer was sequentially removed, providing us with objective measures of the additional impact of this layer on the overall result (*Materials and Methods*, subsection *Assessing omics contributions on subtyping*). We observed (Fig. S2) that bulk RNA, proteomic and metabolomic data emerged as the top contributors to the subtypes, while all layers appeared to be contributing (i.e., small variability was observed across modalities). This further reinforces the notion that different biological components contribute in diverse ways to our understanding of the molecular heterogeneity of AD dementia (*25, 32–34*).

### Cell-type molecular causal drivers of AD dementia subtypes

To further characterize AD dementia subtypes’ disease mechanisms, in a post-hoc analysis we aimed to identify the genes and cells causally driving disease progression in each subtype. For this, we applied the dynamical GENIE3 (dynGENIE3) method (*37*) to each subtype’s snRNA-seq data. DynGENIE3 is a semi-parametric model that describes the temporal changes in gene expression using ordinary differential equations (ODEs), where the transcription function is learned through a non-parametric Random Forest model. Causal regulators for each target cell-type specific gene were determined based on the data-derived variable importance scores from these ODEs. For each subtype-specific dynGENIE3 analysis, we used the participants’ cells-specific genes expression data from snRNAseq (*38*) and their estimated pseudotimes. As described (*Materials and Methods*, *Data*; (*38*)), cells were previously annotated and assigned to seven major cell classes —excitatory neurons (EXC), inhibitory neurons (INH), oligodendrocytes (OLI), oligodendrocyte precursor cells (OPC), astrocytes (AST), immune cells (IMM, including microglia, macrophages, T cells), and vascular and epithelial cells (VASC).

Figures 5A-C present the top 99-percentile causal drivers identified for each AD dementia subtype, ranked by their average direct effect across all cell-types and genes considered. Notably, the primary causal cell-types for the three AD subtrajectories included immune, vascular, and oligodendrocyte precursor cells. Within each subtype, the estimated impacts of these drivers were concentrated around a few cell-type specific genes, with the influence of other cells-genes diminishing rapidly. Numerous genes associated with these three causally leading cell-types (IMM, VASC, OPC) have been previously implicated in the pathogenesis of AD (*39*).

**Figure 5.**
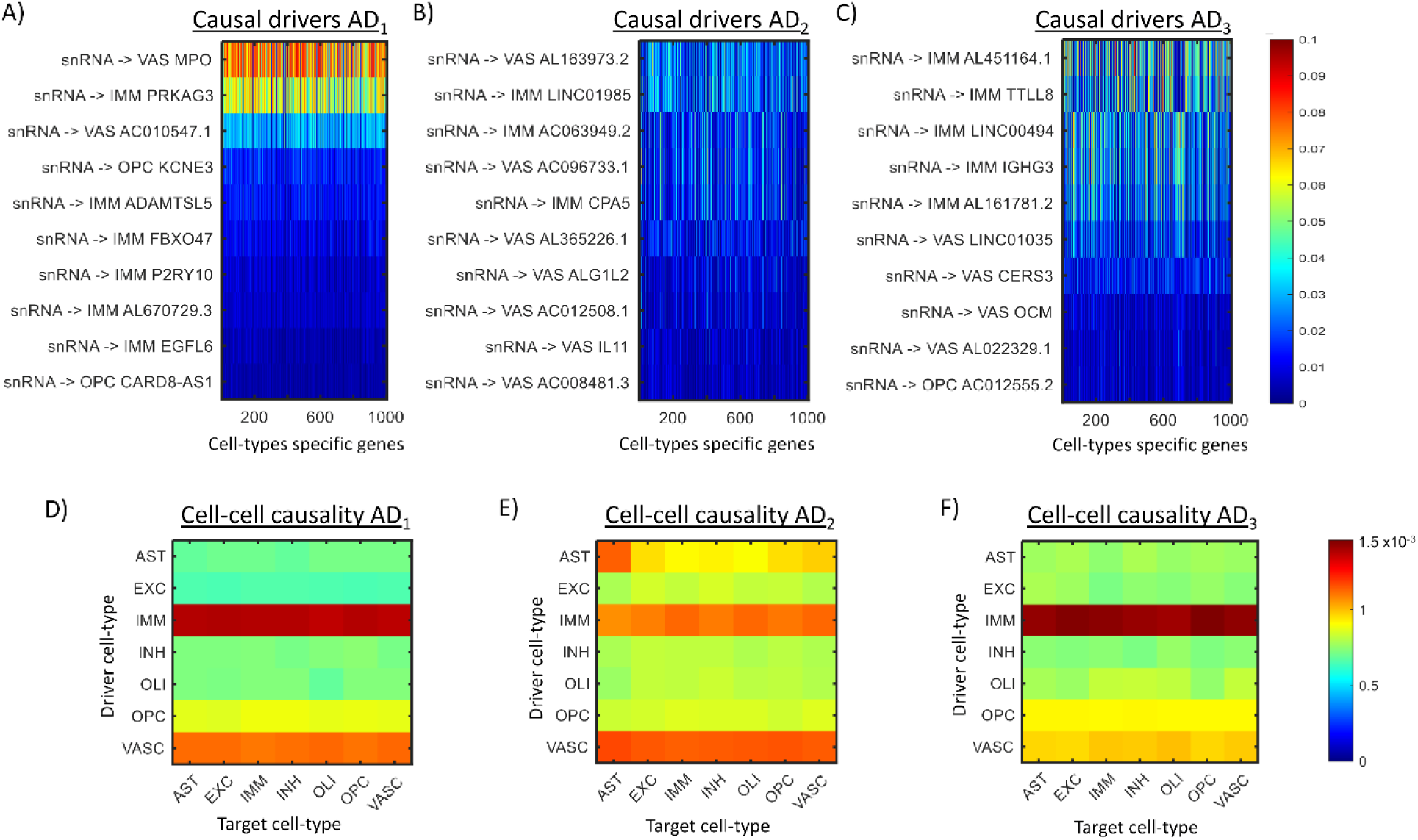
Cellular-level molecular causal drivers for distinct AD subtypes. A) Top 99-percentile causal drivers associated with each subtype. B) Average cell-type to cell-type causal effects.

For AD_1_, the vascular-specific genes MPO (myeloperoxidase) and AC010547.1 have been implicated in blood-brain barrier disruption and cerebrovascular dysfunction in AD (*40, 41*). The MPO enzyme, mainly released by activated neutrophils, is characterized by powerful pro-oxidative and proinflammatory properties (*42*). The immune cell-specific genes PRKAG3, ADAMTSL5, FBXO47, P2RY10, and EGFL6 have been linked to neuroinflammatory processes (*43*). P2RY10 is involved in microglial activation (*43*), which plays a crucial role in AD progression. The OPC genes KCNE3 and CARD8-AS1 have been associated with myelin dysfunction and white matter changes observed in AD (*44*).

For AD_2_, top immune cell-specific genes LINC01985, AC063949.2, and CPA5 may contribute to the neuroinflammatory processes common in AD (*45*). Top vascular-specific genes such as AL163973.2, AC096733.1, AL365226.1, ALG1L2, AC012508.1, and IL11 may also be associated with blood-brain barrier disruption and altered cerebral blood flow observed in AD development (*41*). IL11 has been implicated in neuroinflammation and cognitive decline in AD models (*46*).

AD_3_ is primarily driven by genes associated with immune and vascular cells, with immunoglobulin genes like IGHG3 and IGHM potentially playing a role in neuroinflammation. The altered expression of immunoglobulin genes has been reported in AD brains, suggesting involvement in disease progression (*47*). The vascular-specific gene CERS3 (Ceramide Synthase 3) is part of the ceramide synthesis pathway, known to influence amyloid-β production and accumulation (*48*). OCM (Oncomodulin), another vascular-specific gene, is involved in calcium signaling, which is often dysregulated in AD (*43*).

Furthermore, figures 5D-F present the average causal effects among the different cell-types for each AD subtype. Beyond the top causal drivers (Figs. 5A-C), across all the cell-types and genes, the three subtypes exhibited dominant causal effects stemming from immune, vascular, and astrocytic components. Overall, these findings underscore the complex causal interplay between various cell types in the progression of AD dementia and suggest subtype-specific targets for potential therapeutic interventions.

### Blood, MRI, and psychological traits predictions of brain AD molecular pseudotime

Ultimately, we want the brain data to inform on strategies to prevent and treat AD dementia in living persons. To date, there has been little work translating brain omics pseudotime to living persons (*10*). Here, we leveraged the unique set of paired *ante-mortem* blood omics and psychological risk factors, and *ex vivo* structural neuroimaging (as a proxy for *in vivo* MRI) with the *post-mortem* brain molecular AD dementia pseudotimes. First, we tested the associations between all available blood molecular markers and the pseudotimes. Specifically, for each blood marker, including routine laboratory tests, cell-free DNA, cytokines, monocyte transcripts, proteins, and metabolites, we estimated its Spearman correlation with the pseudotime, adjusting for age, sex, and education. Correlations were standardized by comparison with 1000 randomized permutations (*Materials and Methods, Statistical analyses*), i.e., correcting for potential numerical effects due to sample sizes variations across blood data types and subsequently allowing to compare associations across them.

We identified over a thousand blood markers significantly associated with the brain-derived molecular AD dementia pseudotime (all P<0.05, FDR-corrected). These markers (see Fig. 6A for top 50, and *Table S3* for a complete list) covered a wide range of diverse biological functions, such as lipid metabolism (represented by metabolite TG [51:5]), neurotropic signaling (protein CTF1), cell membrane integrity and signaling (metabolite PC [32:4], protein TIE1), cell adhesion (ITGA11), immune regulation (proteins FCRL2, SDF2L1), inhibitory synaptic function (metabolite Glycine), immune response (protein PIANP), transport within the vascular system (protein PLTP), angiogenesis and vascular function (gene VAV3), cell cycle regulation (protein CSTB), and vascular remodeling (protein TIE1). Significant correlations with the pseudotime were mostly captured by the metabolomic, proteomic, and monocyte transcriptomic markers (Figs. 6A, B). Notably, these results align with the identified cell-type molecular causal drivers in brain for the AD dementia subtypes (Fig. 5, previous subsection), also predominantly linked to broader metabolic, inflammatory, and vascular conditions (*39*).

**Figure 6.**
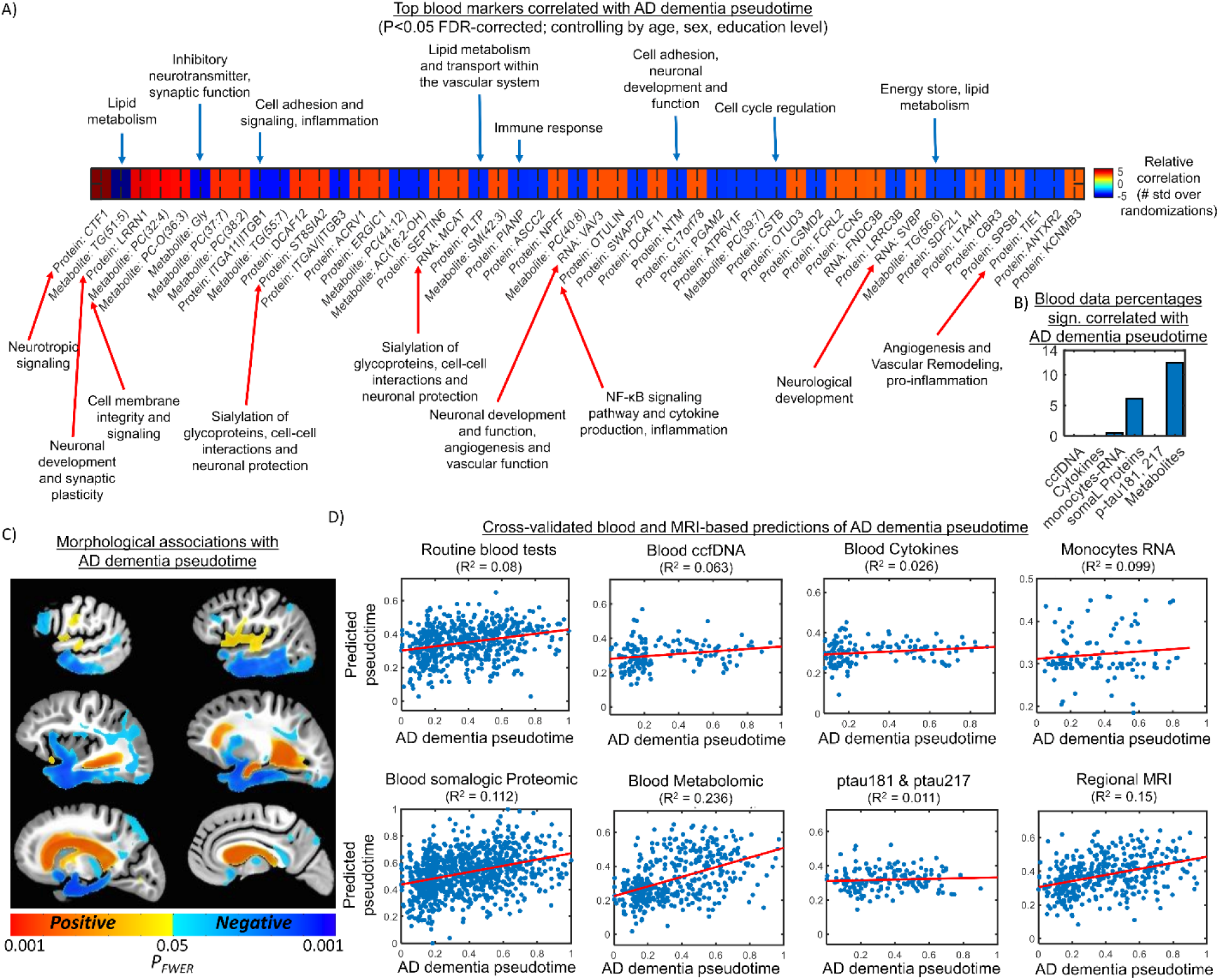
Blood and MRI based predictions of *post-mortem* brain molecular AD dementia pseudotimes. A) Top 50 blood markers significantly correlated with pseudotime (all P<0.05, FDR-corrected; see *Table S3* for a complete list), adjusting for age, sex, and educational level. Correlations were standardized by comparison with 1000 randomized permutations. B) For each blood data modality, percentage of significantly correlated markers with AD dementia pseudotime. C) DBM maps of spatial associations between AD dementia pseudotime and brain structural properties. Warm and cold colors indicate the magnitude of positive and negative associations of regional brain deformation, respectively. D) Blood and regional MRI 10-folds cross-validated predictions of individual AD dementia pseudotime values. See *Materials and Methods, Statistical analyses* and *DBM analysis*.

We used the ex-vivo neuroimaging and deformation-based morphometry (DBM) to characterize brain structural associations with AD dementia pseudotime. We observed (Fig. 6C) that higher pseudotime values were significantly associated (all P<0.05 FWE-corrected) with larger ventricles, as well as less tissue in the temporal lobe and portions of the basal ganglia and parietal lobe. Statistical associations were adjusted by age at death, sex, educational level, postmortem scanning interval and location (see *Materials and Methods*, *DBM analysis*).

In addition, to test whether AD dementia pseudotimes is associated with psychological traits, we ran five separate linear regression models. Outcomes were depressive symptoms, neuroticism, loneliness, conscientiousness, and purpose in life, with pseudotime as predictor, and age at death, sex, and educational level as covariates. Higher pseudotime was positively associated with depressive symptoms (beta=0.60, 95%CI=[0.24, 0.96]; *p* = 0.001), neuroticism (beta= 3.56, 95%CI =[1.83, 5.29]; *p* <0.001), and self-perceived loneliness (beta = 0.38, 95%CI=[0.15, 0.61]; *p* =0.001). Higher pseudotime negatively associated with conscientiousness (beta = -2.79, 95%CI =[-4.57, -1.01]; p = 0.002) and with purpose in life (beta=-0.31, 95%CI =[-0.47, -0.14]; p <0.001).

We next assessed the predictive power of each blood molecular modality, regional MRI metrics, and the psychological traits data to predict AD dementia pseudotime values in new data instances. For each data type, we used several ML regressions with 10-fold cross-validations to predict the individual pseudotime. Regressors included linear regression, support-vector machine (SVM), classification trees, ensembles, neural networks, and Gaussian process regression, adding age, sex and educational level as covariables (*Materials and Methods, Statistical analyses*). Of note, the data modalities were analyzed independently, as the low number of overlapping samples prevented any meaningful combination. The results (Fig. 6D) revealed varying levels of accuracy in blood and regional MRI based predictions of AD dementia pseudotimes, with blood metabolomic and proteomic data showing the strongest predictive capacity (R^2^=0.23 and R^2^=0.11, respectively). Relatively low-dimensional data layers such as cell-free DNA, cytokines, ptau217-ptau181, psychological traits and regional MRI, showed low predictive power (all R^2^ in the range 0.01-0.09), similar to the high-dimensional monocytes RNA data (R^2^=0.09). Overall, these results suggest that while blood and MRI-based biomarkers offer potential for predicting brain-derived AD dementia molecular progression scores in independent samples, with metabolomics demonstrating the greatest promise, several limitations persist. Key challenges include the difficulty in identifying the optimal combination of biomarkers, the use of relatively small sample sizes, the potential influence of unknown confounding variables, and the complexities of biological interactions that may affect the reliability of predictions.

### Blood, MRI, and psychological traits predictions of brain molecular AD dementia subtypes

We continued to leverage the unique set of paired *ante-mortem* blood omics and psychological traits with the *ex vivo* MRI data and *post-mortem* brain multi-omics subtypes. Initial neuroimaging DBM analysis revealed brain structural differences across AD dementia subtypes. As shown in figure 7A, all subtypes exhibited significantly lower temporal lobe volumes and larger ventricles compared to participants with NCI (P<0.05, FWE-corrected). AD_1_ also had significantly smaller volumes in the parietal lobe, while AD_2_ had smaller volumes in several other regions across the brain. When comparing pairs of subtypes, we observed that AD_2_ had significantly larger ventricles relative to AD_3_ (P<0.05, FWE-corrected).

**Figure 7.**
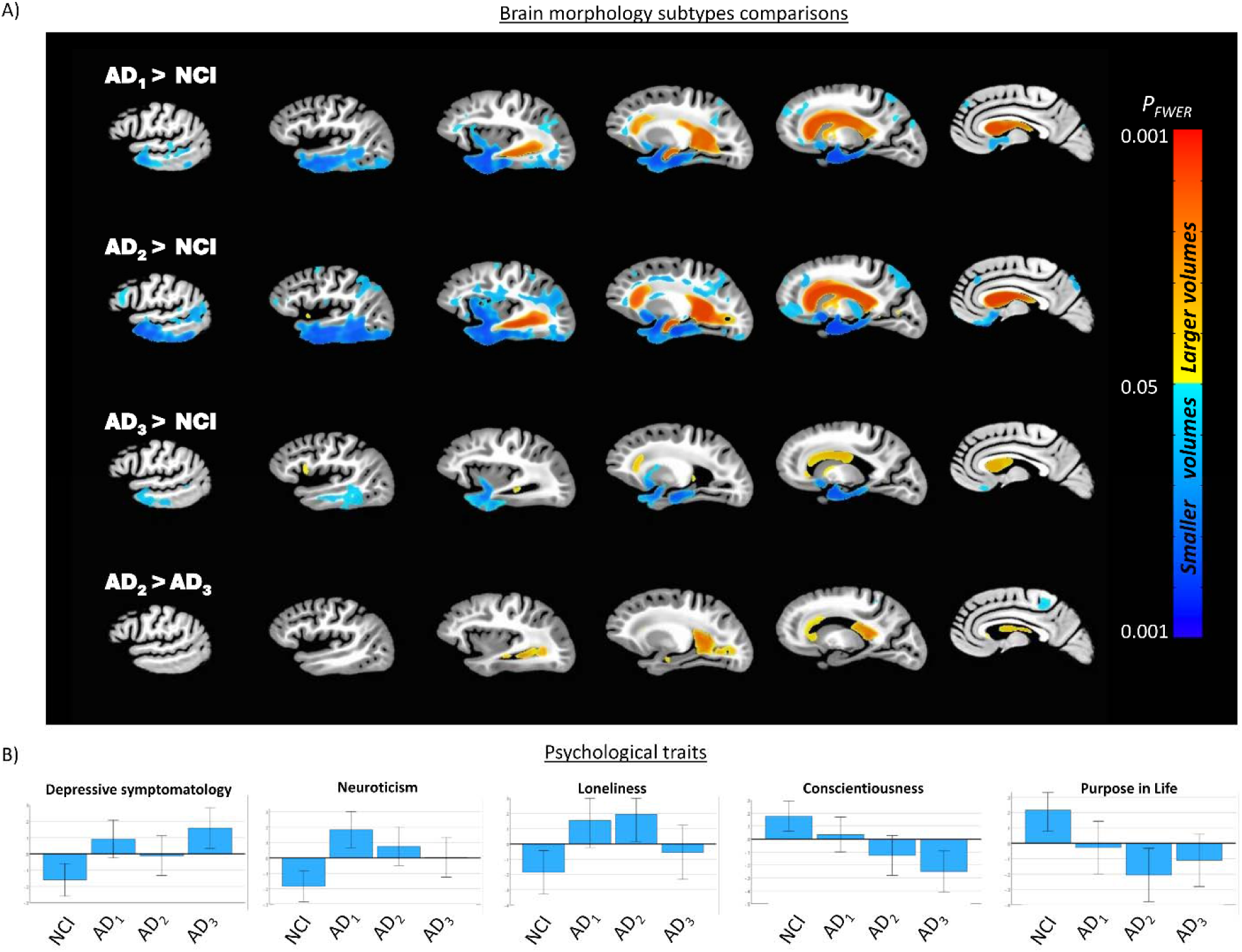
Brain morphology and psychological traits patterns of molecular AD dementia subtypes. A) Structural brain comparisons (all FWE *p*<0.05; including statistical adjustment for age at death, sex, educational level, postmortem scanning interval and location). Warm and cold colors indicate larger and smaller volumes respectively. B) Association of AD dementia subtypes with negative and positive psychological traits (values were z-transformed for illustration purposes; ANCOVA tests included age, sex and educational level as demographic covariates).

Subsequent ANCOVA showed a significant main effect of the AD dementia subtypes on all psychological traits (depressive symptoms: F(3,1173)=6.2, P<0.001; neuroticism: F(3,1129)=7.7, P<0.001; loneliness: F(3,542)=4.7, P=0.003; purpose in life, F(3,541)=5.5, P<0.001; and, conscientiousness, F(3, 802)=7.1, P<0.001) (see *Table S4*). Post-hoc comparisons (Fig. 7B) indicated differential associations between the molecular AD dementia subtypes and the psychological traits relative to the NCI reference group. Mean depressive symptoms in AD_1_ (adjusted mean = 1.5, 95%CI =[1.3, 1.7]) and AD_3_ (adjusted mean = 1.6, 95%CI = [1.4, 1.8]) were different from NCI (adjusted mean = 1.2, 95%CI =[1.02, 1.3]), but AD2 did not differ. Neuroticism and self-perceived loneliness had equivalent differential associations. Mean Neuroticism scores for AD_1_ (adjusted mean = 17.7, 95%CI=[16.9, 18.5]) and AD_2_ (adjusted mean = 17.1, 95%CI=[16.2, 17.8]) were significantly different from NCI (adjusted mean = 15.3, 95%CI =[14.7, 16.1]) by approximately 2 points (Fig. 7B) but AD3 did not differ. Self-perceived loneliness showed the same differential associations as Neuroticism, with AD_1_ (adjusted mean = 2.4, 95%CI=[2.3, 2.6]) and AD_2_ (adjusted mean = 2.5, 95%CI=[2.4, 2.6]) showing significant differences from NCI (adjusted mean = 2.2, 95%CI =[2.2, 2.3]) (Fig. 7B). For positive psychological risk factors, conscientiousness and purpose in life also showed the same differential associations as each other. Mean scores on conscientiousness were lower for both AD_1_ (adjusted mean = 32.5, 95%CI = [31.6, 33.3]) and AD_3_ (adjusted mean = 31.8, 95%CI = [30.9, 32.7]) than NCI (adjusted mean = 34.1, 95%CI = [33.5, 34.8]) by approximately 2 points (Fig. 7B), but did not differ for AD2. Similarly, mean scores on purpose in life were also lower for both AD_2_ (adjusted mean = 3.5, 95%CI = [3.4, 3.5]) and AD_3_ (adjusted mean = 3.5, 95%CI = [3.4, 3.6]) than NCI (adjusted mean = 3.6, 95%CI = [3.6, 3.7]), but not AD1. Finally, when comparing subtype pairs, we observed significant differences in conscientiousness (P=0.045) between AD_1_ (adjusted mean = 33.4, 95%CI=[32.6, 34.1]) and AD_3_ (adjusted mean = 31.8, 95%CI=[30.9, 32.7]). In sum, as indicated by our results, neuroticism and self-perceived loneliness shared a similar profile for AD_1_ and AD_2_ vs NCI, while conscientiousness and purpose in life presented a similar profile for AD_2_ and AD_3_. In line with the observed cognitive patterns (Fig. 4E), AD_2_ consistently had more differences with NCI than the other two subtypes, with four traits (higher neuroticism, high self-perceived loneliness, lower purpose in life, and lower conscientiousness) being significantly different.

We then evaluated the ability of each blood molecular modality, the regional MRI metrics, and the psychological traits to predict the AD dementia subtypes in new data instances. For each data type, we used several ML classifiers with 5-fold cross-validations to predict the individually assigned subtypes in AD participants (i.e., categories AD_1_, AD_2_, or AD_3_), by analyzing each versus NCI or the other subtypes. Classifiers included support-vector machine, classification trees, naïve Bayes, neural networks, *k*-nearest neighbor, linear discriminative analysis and ensembles, adding age, sex and educational level as covariables (*Materials and Methods, Statistical analyses*). The results (Fig. 8) revealed varying levels of area under the ROC curve (AUC) in blood, regional-MRI and psychological traits-based predictions of AD subtypes. Low-dimensional data layers such as cell-free DNA, cytokines, and ptau217-ptau181 showcased relatively high accuracy in distinguishing AD subtypes from NCI. However, their ability to differentiate between specific subtypes (AD_1_ vs AD_2_, AD_1_ vs AD_3_, and AD_2_ vs AD_3_) was significantly lower, except for AD_1_ vs AD_3_ predictions (reaching 0.66 to 0.71 AUC values). Contrary, psychological traits demonstrated a limited capacity to predict AD subtypes vs NCI, but exhibited the strongest predictive capability among all data modalities to distinguish AD_2_ vs AD_3_ and AD_1_ vs AD_3_, reaching 0.83 and 0.76 AUC values, respectively. Monocytes RNA exhibited the lowest overall accuracy, with limited success in predicting AD dementia subtypes from NCI and in differentiating between pairs of subtypes. Proteomics and regional-MRI demonstrated moderate levels of AUC for both tasks (reaching from 0.68 to 0.86, and from 0.58 to 0.73 AUCs, respectively). Blood metabolomics, however, emerged as the most promising omic in blood, achieving the highest accuracy in predicting AD subtypes from NCI (AUC= 0.99) and exhibiting medium capacity in differentiating between specific subtypes (from 0.62 to 0.68 AUC values). Overall, these results are consistent with the predictions for AD dementia pseudotimes, suggesting that blood molecular, neuroimaging and psychological risk factors data may be valuable for predicting brain-derived AD dementia taxonomies. Among these, metabolomics demonstrates the greatest potential to distinguish AD vs NCI, while psychological traits showed the strongest capacity to differentiate among distinctive molecular dementia subtypes in AD. However, significant limitations remain, including the limited sample sizes and optimal markers selection, highlighting the need for substantial enhancement.

**Figure 8.**
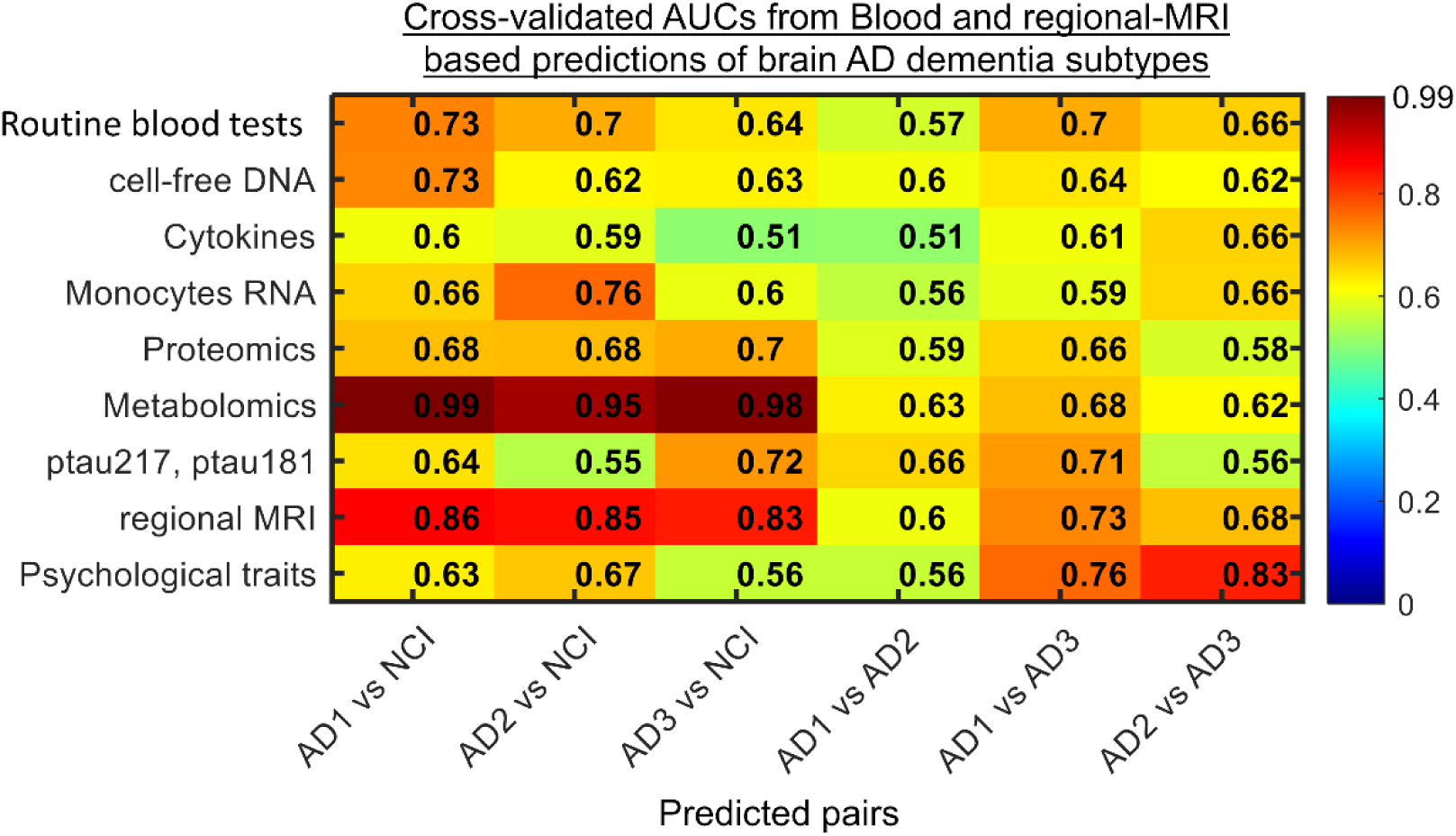
Blood, regional MRI and psychological traits-based predictions of *post-mortem* brain multi-omics AD dementia subtypes. Prediction AUC for every pair of subgroups included in the classification tasks. Notice that the first three columns correspond to subtype-specific vs NCI classifications, while the last three columns correspond to subtype vs subtype classifications.

## DISCUSSION

This study represents a significant advancement in the characterization of AD dementia by taking the first steps towards successfully translating a robust multi-omics molecular taxonomy derived from *post-mortem* brain samples into living individuals. By mapping the substantial heterogeneity of AD dementia into distinct brain molecular subtrajectories, we identified unique cellular and molecular drivers, along with different pathological, clinical, morphological and psychological risk factor profiles. Moreover, we established essential groundwork for translating the molecular subtrajectories to data generated from living humans. The implications of our findings extend far beyond mere classification of AD; they open promising avenues for the development and implementation of personalized interventions tailored to individual molecular profiles, aiming to enhance therapeutic outcomes for a range of brain diseases and resilience to those diseases.

Molecular diagnostics have played a pivotal role in advancing precision medicine for cancer (*49, 50*). Building on these achievements, molecular-based pseudo-temporal studies in AD aim to unravel the temporal molecular changes during its long-term and varied progression. By applying manifold learning techniques to RNA-Seq data from postmortem brain tissues (*4, 7, 51*), early studies have estimated AD pseudotimes that correlate with the severity of neuropathologies and cognitive performance. These models also have shed light on the key transcripts and pathways implicated in AD progression. Nevertheless, because these studies rely mostly on bulk RNA data, they often overlook crucial factors such as epigenetic modifications, protein alterations, metabolic dysfunction, neuroinflammation, and mitochondrial abnormalities that interact with gene function in AD. Furthermore, they may not fully capture the cellular context, intercellular signaling, and diversity inherent to AD heterogeneity. To account for this, we recently integrated four molecular omics, subsequently identifying distinct, molecularly differentiated, and statistically stable pseudo-temporal AD subtrajectories, and measuring individual-specific progression along these (*10*).

By aggregating eight complementary molecular layers, including DNA methylation, RNA sequencing (single-nucleus and bulk), proteomics (tandem mass tag, glycated, and glycosylated), metabolomics, and lipidomics, here we have achieved a more comprehensive characterization of AD dementia progression than previous studies using fewer omics (*3, 7, 10, 51, 52*). The refined AD dementia pseudotime offers notably better predictions of cognitive performance and decline versus our earlier four-omics model (*10*), highlighting the added value of incorporating more molecular data. Additionally, in the three identified molecular subtypes of AD dementia, the detection of cell-type specific molecular drivers emphasizes the causal roles of immune, vascular, and oligodendrocyte precursor cells in AD pathogenesis, in alignment with recent findings (*39*). The unique features of each subtype —distinctive molecular contributors, brain morphologies, sex distributions, cognitive and psychological risk profiles— stress the importance of personalized therapeutic approaches considering molecular, demographic and psychological factors. Crucially, our developed predictive models were able to successfully convert advanced brain-derived molecular profiles —AD dementia pseudotimes and subtypes— into blood-based markers, MRI metrics, and psychological traits, providing the first-generation translatable biomarker signatures for distinct AD molecular pathways. While promising, these models also present opportunities for refinement. We aim to enlarge sample sizes and increase overlap between *ante-mortem* and *post-mortem* samples. Furthermore, the ability to generate *ante-mortem* signatures of brain molecular subtrajectories, when paired with induced organoids from the same individuals, has the potential to revolutionize the approach to precision medicine for AD and other brain diseases. Indeed, we have shown that we can model several AD/ADRD clinical and pathologic traits from induced neurons and astrocytes from these participants (*53–56*). This suggests that we may be able to do the same for the molecular subtrajectories.

We note the strengths of the study ROSMAP are community-based with extraordinarily high follow-up and autopsy rates limiting bias due to selective attrition. The paired survey, clinical, blood, imaging, neuropathological, and multi-level deep brain omics data in large numbers of the same humans is not available in any other extant study. Nevertheless, several limitations should be addressed in future research. Validation in larger, diverse cohorts is necessary to confirm the robustness and generalizability of the identified AD dementia pseudotimes/subtypes and the translational models. As noted, no other study in AMP-AD or similar data repository presents all *post-mortem* and *in-vivo* data types used in this study, hindering replication in independent samples. To partly overcome this limitation, we performed extensive cross-validation tests in our translational analysis, evaluating portability of predictions. Further investigation into the causal relationships between the identified subtype-specific cellular drivers of AD dementia progression could provide precise therapeutic targets, which will require further experimental testing and validation. The translation of brain-derived molecular profiles to blood, MRI and psychological-based markers, while promising, requires further optimization to improve predictive accuracy (e.g., increasing sample sizes and features selection for high dimensional modalities). Furthermore, while considering both sexes, our findings will be limited to ROSMAP’s predominantly White non-Latinx educated American population. The Rush Alzheimer’s Disease Center (RADC)’s recent developments in profiling Black Americans (*57*) is promising, including more than 200 autopsies to date and continuing data generation for future analyses. For brain omics, RADC is also building a similar multi-omics brain platform in a very large number of Black/Mixed and White Brazilians across the entire age and education spectrum. Thus, future extension of this study’s findings to racially diverse populations will soon be feasible.

## MATERIALS AND METHODS

### Data

#### Ethics statement

The Religious Orders Study (ROS) and the Rush Memory and Aging Project Study (MAP) (*58*) were each approved by an Institutional Review Board of Rush University Medical Center. All participants enrolled without known dementia and signed an informed consent and Anatomical Gift Act agreeing to annual detailed clinical evaluation and *post-mortem* brain donation; in addition, they signed a repository consent allowing their data to be shared. Data documentation and sharing documents can be obtained at https://www.radc.rush.edu. The study was conducted according to Good Clinical Practice guidelines, the Declaration of Helsinki, and Institutional Review Boards (adni.loni.usc.edu). Study subjects and/or authorized representatives gave written informed consent at the time of enrollment for sample collection and completed questionnaires approved by each participating site Institutional Review Board (IRB).

#### Data origin

This study used brain and blood multi-omics molecular, neuropathological, and/or clinical data from ROS and MAP (*58*) cohorts (N_total_=1,189; see *Table S1* for demographic characteristics).

The *post-mortem* brain multi-omics data included DNA methylation, single-nucleus and bulk RNA-seq expression, proteomic, glycoproteomic, metabolomic and/or lipidomic from the dorsolateral prefrontal cortex (DLPFC) of 1,515 autopsied brains. Only 1,189 participants with at least 2 molecular data modalities generated from brain tissue were considered in subsequent analyses (see Fig. S1A). All these data were generated in previous studies, as described in (*59–64*), and most is available at the Accelerating Medicines Partnership Alzheimer’s Disease knowledge portal (AMP-AD; www.synapse.org). The following Synapse IDs were used: syn23650894, syn3388564, syn17015098, syn26007830, syn22024496. The metabolomic data was generated by the Alzheimer’s Disease Metabolomics Consortium (ADMC; see also Acknowledgments). Epigenome-wide 5mC data was generated using the Infinium MethylationEPIC v2.0 BeadChip (*Illumina*, https://www.illumina.com/products/by-type/microarray-kits/infinium-methylation-epic.h). High-quality reads were mapped to the GRCh38 human genome. After data pre-processing, removal of technical batch variance, and multi-steps quality control (QC), a total of 885,040 CpGs were obtained. Single-nucleus RNA-seq (Synapse ID syn23650894) was performed across 2.3 million isolated nuclei, and cells were previously annotated and assigned to seven major cell classes —excitatory neurons (EXC), inhibitory neurons (INH), oligodendrocytes (OLI), oligodendrocyte precursor cells (OPC), astrocytes (AST), immune cells (IMM, including microglia, macrophages, T cells), and vascular and epithelial cells (VASC) — as described (*38, 65*). Bulk RNA-seq (syn3388564) generated using next-generation sequencing (NGS) was processed by estimating gene counts with Kallisto and normalizing as described (*66*). Proteomic data (syn17015098) was generated for a total of 8,356 protein levels quantified using tandem mass tags (TMT) as described (*67*). Glycoproteomic data included 11,012 glycopeptiforms quantified using liquid chromatography with tandem mass spectrometry as described (*68*). Metabolon-wide data (syn26007830) was generated on the *Metabolon Precision Metabolomics platform* (Metabolon, Inc., Morrisville, USA) as described (*69*). Lipidomic data was generated using non-targeted mass spectrometry as described (*70*), obtaining a broad coverage of the brain lipidome, with for 346 lipids.

The *in-vivo* blood omics included (see Fig. S1B) cell-free DNA (6 markers), monocytes RNA-seq (52,650 transcripts; syn22024496), SomaLogic proteins (7,289), p-tau181 and p-tau217, cytokines (4), and/or Biocrates metabolites (292) for a subset of 1,108 ROSMAP participants with brain multi-omics data (Fig. S1). Data were generated in previous studies (*71–73*).

Annual administration of cognitive tests were incorporated into summary measures of 5 domains of cognitive function (episodic memory, visuospatial ability, perceptual speed, semantic memory and working memory (*74–77*)), and a global cognition measure computed by averaging the 5 summary scores. In addition, for each cognitive domain, the person-specific random slope was estimated as the rate of change in the variable over time. It comes from linear mixed effects model with annual variable as the longitudinal outcome. The model controls for age at baseline, sex, and years of education (*76, 78*). Assignment of NCI, MCI or AD dementia categories was performed in correspondence with most likely clinical diagnosis at the time of death. All available clinical data were reviewed by a neurologist with expertise in dementia, and a summary diagnostic opinion was rendered regarding the most likely clinical diagnosis (*16–18*). All subjects underwent *post-mortem* neuropathologic evaluations, including uniform structured assessment of AD pathology, cerebral infarcts, Lewy body disease, TDP-43 cytoplasmatic inclusions in neurons and glia, and other pathologies common in aging and dementia (*14, 15*). The pathologic diagnosis of AD uses NIA-Reagan and modified CERAD criteria, and the staging of neurofibrillary pathology uses Braak Staging (*12, 13*).

Depressive symptoms were assessed annually using a 10-item form (*21*) of the Center of Epidemiologic Study-Depression Scale (CES-D; (*22*)). Because we did not observe any systematic change in CES-D scores in participants during follow-up (*23*), we used each participant’s average CES-D score across all evaluations to enhance our assessment of the enduring disposition to experience depressive symptoms. Neuroticism was assessed at baseline using either 12 (n=1031) or 6 items (n=105) from the NEO Five-Factor Inventory (*79*), which was administered at baseline or near baseline as previously reported (*80*). Loneliness was assessed at baseline in MAP participants only, using a 5-item version from a modified scale of the de Jong-Gierveld Loneliness scale (*24, 81*). Conscientiousness was assessed at baseline using 12 items from the NEO Five-Factor Inventory as previously described (*82*). Purpose in life was assessed at baseline in MAP participants only, using a 10-item scale derived from Ryff’s Scales of Psychological Wellbeing as previously described (*20, 83*).

#### Data preprocessing

Before applying the *mcTI* approach, each molecular feature’s quality-controlled values (e.g., CpG site methylation level, gene abundance, protein, glycoprotein, metabolite or lipid concentration) were adjusted for relevant covariates using robust additive linear models. Covariables included age, sex and educational level for both *in-vivo* and *post-mortem* data, plus *post-mortem* interval (PMI) in hours, sample pH, RNA integrity number (RIN) and batch number when applicable.

#### Multimodal contrastive Trajectories Inference (mcTI) definition

Initially proposed in (*10*), the mcTI algorithm has been recently updated to improve specific aspects. Given a set of multiple ‘omic’ data types, this method provides two estimations for each participant: 1) molecular disease pseudotime, e.g., reflecting how close (in terms of multi-level molecular alterations) each participant is to developing AD dementia, and 2) a putative disease subtype, corresponding to a distinctive disease subtrajectory that the participant aligns with. Initially, each data modality’s high number of features/markers is reduced by identifying the low-dimensional pattern enriched in the target subpopulation (here, subjects diagnosed with AD dementia) relative to the reference/background subpopulation (here, NCI).

First, a contrastive Principal Component Analysis (cPCA (*84*)) is performed, reducing each data modality to a few components capturing the AD dementia associated patterns (*52*). The contrasted principal components (cPC) resulting from all different data modalities are subsequently used as input in a probabilistic Principal Component Analysis (pPCA), which identifies fused contrasted components capturing the common variance across all modalities’ contrasted/enriched patterns, while dealing with missing data, i.e., varying N’s for each omic layer. A subject-subject dissimilarity network (SDN) is then constructed based on the inter-individual Euclidian distances across the resulting fused cPCs. This SDN is used to calculate the shortest path from any participant to the background group. Each shortest path is defined as the concatenation of relatively similar subjects in the integrated multi-omics molecular space that minimizes the distance to the background’s centroid. The position of each subject in the corresponding shortest path reflects the individual distance to the background subpopulation (NCI) or, if analyzed in the reverse direction, to the undesired state (AD dementia). Thus, to quantify the distance to these two extremes (NCI or AD dementia), the individual multi-omics pseudotime is calculated as the shortest distance value to the background’s centroid, relative to the maximum population value (i.e., values are standardized between 0 and 1). Relatively low or high values indicate greater or lesser distance on the path to develop AD dementia.

Once the pseudotimes are estimated, before proceeding to identify subtypes, all the fused contrasted components are statistically adjusted by the pseudotime values via robust regression. These new disease-adjusted components are then non-linearly embedded into a two- or three-dimensional space via t-SNE (*85*), where the participants are subsequently clustered. The number of t-SNE dimensions is selected to maximize results stability across perplexity values (i.e., in the range [5 to 50] with 5 as step size) and clustering criteria. The resulting clusters will define the final putative subtypes, i.e., after adjusting the individually embedded data for disease progression as captured by the pseudotime. To identify the optimum number of clusters/subtypes, a majority rule across the Calinski–Harabasz and Silhouette criteria is used (*35, 36*). The subtypes’ stability and significance are evaluated via randomized permutations. Following the method proposed in (*3*), subtype stability is defined as the rate at which pairs of subjects group together into the same subtypes upon repeated clustering on random subsets of the input data. Analyses were performed in MATLAB version R2021b.

#### Assessing markers contributions on pseudotime

For each dataset and molecular omics modality, the total contribution *C_i_* of each modality-specific marker *i* to the obtained reduced representation space (and the multi-omics pseudotime) was quantified as (*10*):

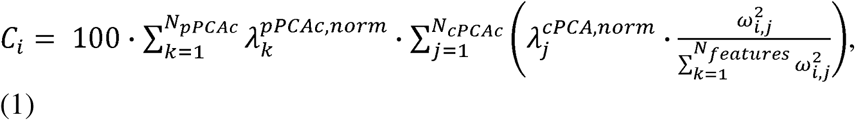

where 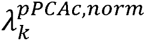 is the normalized eigenvalue of the pPCA component *k* from a total of *N_pPCAc_*resulting components after fusing all data modalities, *N_cPCAc_* is the number of contrasted principal components for marker *i*’s corresponding data modality, 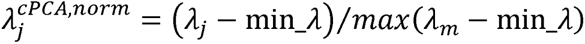 is the normalized eigenvalue of the contrasted principal component *j*, min_λ is the minimum obtained eigenvalue across all markers for *j*, *ω_i,j_* is the loading/weight of the marker *i* on the component *j*, and *N_markers_* is the total number of modality-specific markers in *i*’s corresponding data modality.

#### Assessing omics contributions on subtyping

For each molecular omics modality *i*, its contribution to the obtained subtypes was calculated as:

(i) *i* was removed from the *mcTI* algorithm’s input data, and a new set of AD subtypes was obtained (i.e., without any information from *i*),
(ii) an index reflecting the level of dependence in modality *i* for obtaining the original AD subtypes: 1-*nM1*, where *nM1* is the *normalized Mutual Information* between the two obtained AD subtyping configurations (i.e., with and without considering modality *i*). Notice that a value of 1 would imply a high level of dependency in modality *i*, and 0 otherwise.

The above indices were normalized to express percentages across the molecular modalities.

#### Identification of Subtypes Signatures

Our objective was to identify distinct molecular alterations across each subtype by detecting omic features that significantly deviated from typical values in the NCI reference group (P<0.05, FDR-corrected), while remaining statistically stable in other subtypes. For each subtype *i* and omic feature *j*, we employed an ANCOVA test with 500 randomized permutations to calculate their alteration levels, generating (F*_i,j_*and P*_i,j_*) statistics, adjusted for age, sex, and educational level. We then identified all omic features that were significantly different for any subtype (P<0.05, FDR-corrected). From this pool, we selected the omic features that were significantly altered for subtype *i* but non-significantly altered for the other subtypes. These selected omic features constituted the distinct molecular signature for subtype *i* (see Files S1-S3).

For interpretation, within subtype *i*, the selected features were also ranked according to their propensity to be predominantly different for *i* while being preserved in the other subtypes. To quantify this tendency, we calculated an ’anisotropy’ index, defined as follows (*86, 87*): 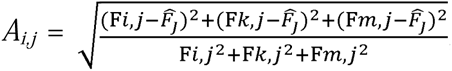, where *k* and *m* represent the other subtypes and 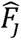 is the average F-value for feature *j* across subtypes. This formula can accommodate different numbers of subtypes by including or excluding terms accordingly. The index (*A_i,j_*) yields a scalar value between zero and one, indicating the distinctiveness of feature *j* concerning subtype *i*; values closer to one suggest a stronger distinctiveness. This approach is an extension of the concept of fractional anisotropy commonly employed in diffusion neuroimaging (*86, 87*) and relates to the eccentricity of conic sections in three dimensions, normalized to a unit range. Notice that while F-values may not be directly comparable across data modalities due to varying sample sizes, the *A_i,j_* values allow for meaningful comparisons since they are standardized across subtypes.

#### Cell-type specific causality analysis

To identify genes specific to cell types that causally drive disease progression within each subtype, we employed the dynamical GENIE3 (dynGENIE3) method (*37*) to analyze the corresponding snRNA-seq data. DynGENIE3 (available at https://github.com/vahuynh/dynGENIE3), is a semi-parametric approach that models temporal changes in gene expression using ordinary differential equations (ODEs). This method determines causal regulators for each target cell-type specific gene based on data-derived variable importance scores. For our subtype-specific analyses, we utilized the participants’ gene expression values specific to cell types, ordering the participants by their estimated AD dementia pseudotimes. We utilized Random Forest trees to optimize the ODEs and performed five-fold partitions of each subgroup of participants, thereby simulating different experimental pools and enhancing the robustness of our estimates. DynGENIE3 generated subtype-specific gene regulatory networks, providing numerical values that reflect the direct impact of each cell-type specific gene on other genes across all considered cell types.

#### DBM analysis

In a subset of 599 (191 NCI, 145 AD_1_, 149 AD_2_, 114 AD_3_) decedents with postmortem neuroimaging, we examined the associations of pseudotime and AD subtype with brain morphology. Briefly, after one month postmortem, the cerebral hemisphere selected for neuropathological examination was first imaged with T2-weighted sequences on one of four 3-Tesla scanners, as previously reported (*11*). These T2 images were later corrected for N4 bias field inhomogeneity (*88*) and non-linearly registered to an in-house postmortem hemispheric template (*89*) using Advanced Normalization Tools (ANTs)(*90*). This registration was used to compute voxel-wise Jacobian determinant maps, which we then log-transformed and smoothed with a 4mm FWHM Gaussian kernel. Higher values in these deformation maps indicated large volumes that were contracted to fit the template, whereas lower values indicated smaller volumes that were expanded to the template. For this analysis, we implemented two general linear models using FSL PALM (*91, 92*), which could assume different means and variances across scanners. In the first model, we entered AD dementia pseudotime as the explanatory variable (EV) of regional deformation and controlled for age at death, sex, years of education, *post-mortem* scanning interval, and scanner location. In the second model, we entered four indicator variables for AD dementia subtypes as EVs of interest, also adjusted for demographics and postmortem conditions, and ran contrasts for each group pair. P-values were computed from 500 permutations using tail approximation (*93*), threshold-free cluster enhancement, and family-wise error (FWE) correction. Associations with FWE p≤.05 were considered statistically significant.

#### Additional statistical analyses

Cognitive, neuropathological and blood correlations with brain-derived AD dementia pseudotimes were estimated via partial Pearson correlations (adjusting by age, sex, and education) and standardized by comparison with 1000 randomized permutations. Clinical-level associations with AD dementia pseudotimes were estimated via an ANCOVA analysis (with clinical diagnosis as grouping variable, adjusting by age, sex, and education) and 1000 randomized permutations. Blood, regional MRI and psychological data-based machine-learning predictions of AD dementia pseudotimes used linear regression, SVM, classification trees, ensembles, neural networks, and Gaussian process regression, under a 10-fold cross-validation scheme. Classifications of AD subtypes were based on SVM, classification trees, naïve Bayes, neural networks, *k*-nearest neighbor, linear discriminative analysis and ensembles, under a 5-fold cross-validation scheme and restricted to AD and NCI participants. Both regression- and classification-based predictions included age at death, sex and educational level as covariables.

## Supplementary Materials

**Figure S1** | UpSet graphs with available data modalities.

**Figure S2** | Contribution of brain omics to AD dementia molecular subtypes.

**Table S1** | Main demographic and data characteristics.

**Table S2** | Routine lab blood tests, cell-free DNA, cytokines and regional-MRI markers.

**Table S3** | Blood markers significantly associated with the brain-derived molecular AD dementia pseudotime.

**Table S4** | Post-hoc subtype-subtype comparisons on neuroticism, conscientiousness, depressive symptoms, loneliness, and purpose in life.

**File S1** | Distinctive multi-omics signature AD_1_.

**File S2** | Distinctive multi-omics signature AD_2_.

**File S3** | Distinctive multi-omics signature AD_3_.

## Data and materials availability

All data needed to evaluate the conclusions in the paper are present in the paper and/or the Supplementary Materials. ROSMAP molecular data are available for general research at the AMP-AD knowledge portal (https://www.synapse.org/), according to the following requirements for data access and attribution (https://adknowledgeportal.synapse.org/DataAccess/Instructions). Detailed ROSMAP neuropathological, neuroimaging, psychological and clinical data are available at the RADC Research Resource Sharing Hub (<u>radc.rush.edu</u>), pending scientific review and a completed material transfer agreement (see <u>radc.rush.edu/requests.htm</u>). The used multi-omics stratification method (*mcTI*) is freely available as part of the *NeuroPM-box* software (*94*) (<u>neuropm-</u> <u>lab.com/neuropm-box.html)</u>, which updated codes regularly released.

## Acknowledgments

We appreciate the participants of ROS and MAP for the time generously given for data collection and for agreement to brain donation. We also acknowledge staff of Rush Alzheimer’s Disease Center for data collection and management. This project was partly supported by the following awards to YIM: Canada Research Chair tier-2, CIHR Project Grant 2020, and Weston Family Foundation’s Transformational Research in AD 2020. We used the computational infrastructure of the McConnell Brain Imaging Center, supported in part by the *Brain Canada Foundation*. In addition, this work was supported by National Institutes of Health grant: R01AG17917, P30AG10161, P30AG72975, R01AG015819, U01AG61356, U01 NS100599, R01 AG064233, RF1 NS139975, P30 AG072975. We would also like to thank the Paul M. Angell Family Foundation for their generous financial support given towards this work, and Michael Urbut for his support.

