## Supplementary information for "Translating the Post-Mortem Brain Multi-Omics Molecular Taxonomy of Alzheimer’s Dementia to Living Humans"

Yasser Iturria-Medina et al.

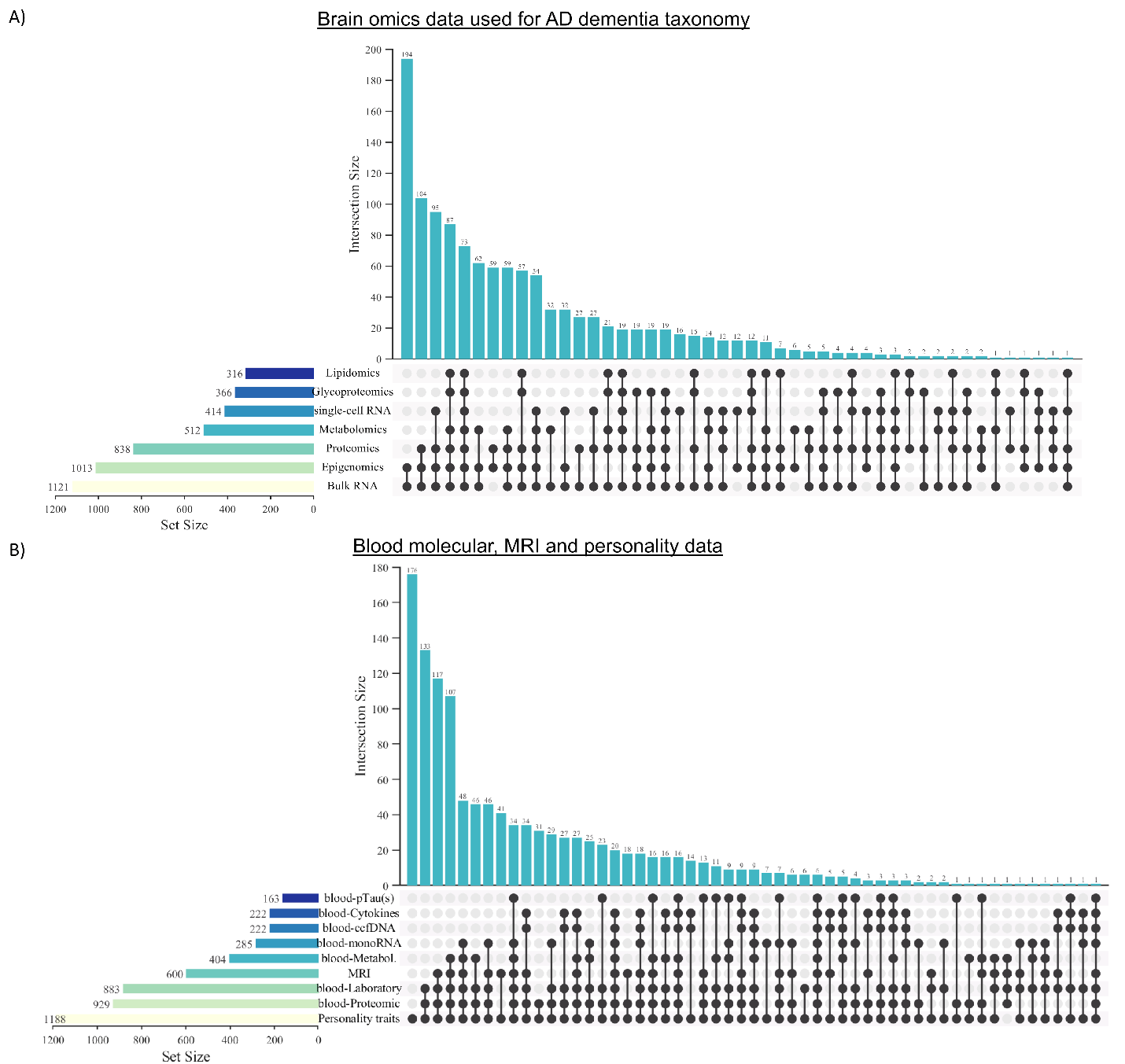

**Figure S1 |** UpSet graphs for available data modalities. A) Post-mortem brain omics used for AD dementia taxonomy. B) Blood, MRI and personality data used for cross-validated predictions of AD dementia pseudotimes and subtypes.

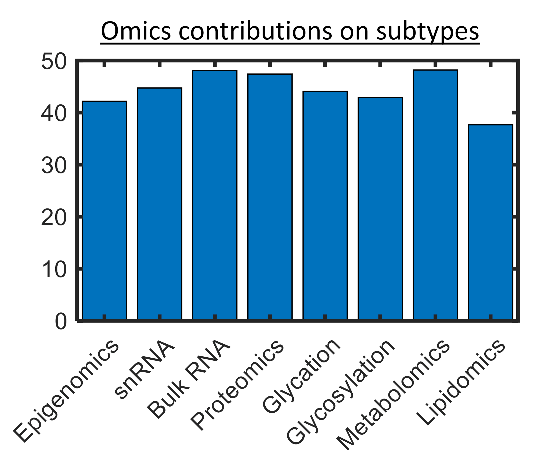

**Figure S2 |** Contribution of brain omics to AD dementia molecular subtypes. Values are percentages. For technical details, see *Materials and Methods*, *Assessing data contributions on subtrajectories*.

**Table S1 |** Main demographic and data characteristics. Data are number n (%) or Mean (SD).

|  | Mean (SD) or % (n) | | | | |
| --- | --- | --- | --- | --- | --- |
| **Characteristics** | Whole sample | NCI (background) | AD_1_ | AD_2_ | AD_3_ |
| N | 1189 | 390 | 294 | 255 | 249 |
| Age at baseline, mean years, (SD) | 80.7 (6.9) | 78.9 (7.1) | 82.3 (6.7) | 81.0 (6.8) | 80.7 (6.9) |
| Age at death, mean years (SD) | 89.6 (6.5) | 87.6 (6.8) | 91.6 (6.2) | 90.6 (5.7) | 89.5 (6.2) |
| Female, n (%) | 378 (32) | 260 (67) | 247 (84) | 178 (70) | 125 (50) |
| Educational attainment, mean years (SD) | 16.1 (3.6) | 16.2 (3.6) | 15.9 (3.7) | 15.7 (3.3) | 16.7 (3.6) |
| Neuroticism, baseline, (SD) | 16.5 (6.5) | 15.5 (6.6) | 17.7 (6.4) | 16.9 (6.3) | 16.3 (6.4) |
| Depressive symptoms, mean score (SD) | 1.2 (1.7) | 1.1 (1.5) | 1.4 (1.8) | 1.2 (1.6) | 1.3 (1.7) |
| Loneliness, baseline (SD) | 2.4 (0.6) | 2.2 (0.6) | 2.5 (0.6) | 2.5 (0.6) | 2.3 (0.6) |
| Purpose in life, baseline (SD) | 3.5 (0.5) | 3.7 (0.5) | 3.5 (0.4) | 3.4 (0.4) | 3.5 (0.5) |
| Conscientiousness (SD) | 33.2 (5.5) | 34.0 (5.4) | 33.5 (5.1) | 32.4 (4.8) | 31.8 (6.7) |
| Pseudotime, mean (SD) | 0.4 (0.2) | 0.1 (0.1) | 0.5 (0.2) | 0.5 (0.2) | 0.5 (0.2) |

**Table S2 |** Routine lab blood tests, cell-free DNA, cytokines and regional MRI markers.

| **Data type** | **Names** |
| --- | --- |
| Routine laboratory tests | Anemia Blood urea nitrogen (BUN) Calcium level Carbon dioxide level Chloride level Cholesterol/HDL ratio Creatinine level Estimated glomerular filtration rate Fasting status Glucose level Hematocrit Hemoglobin Hemoglobin A1c level HDL cholesterol level LDL cholesterol level Mean corpuscular hemoglobin (MCH) Mean corpuscular hemoglobin concentration (MCHC) Mean corpuscular volume (MCV) Platelet count Potassium level RBC count RBC distribution width Red blood cell distribution width in blood samples Sodium level TSH level Total cholesterol level Triglyceride level WBC count |
| cell-free DNA | Short, med, longf, nd6, gdnaned, gdnahex |
| Cytokines | crp_ug, tnfr1, il6, tnfa |
| regional MRI | R2_roi39, R2_roi26, R2_roi42, R2_roi43, R2_roi33, R2_roi15, R2_roi02, R2_roi05, R2_roi41, R2_roi01, R2_roi18, R2_roi09, R2_roi25, R2_roi20, R2_roi29, R2_roi22, R2_roi04, R2_roi40, R2_roi37, R2_roi19, R2_roi16, R2_roi17, R2_roi38, R2_roi36, R2_roi13, R2_roi03, R2_roi12, R2_roi11, R2_roi34, R2_roi06, R2_roi21, R2_roi08, R2_roi24, R2_roi10, R2_roi14, R2_roi30, R2_roi32, R2_roi07, R2_roi28, R2_roi23, R2_roi31, R2_roi27  VOLUME_roi39, VOLUME_roi26, VOLUME_roi42, VOLUME_roi43, VOLUME_roi33, VOLUME_roi15, VOLUME_roi02, VOLUME_roi05, VOLUME_roi41, VOLUME_roi01, VOLUME_roi18, VOLUME_roi09, VOLUME_roi25, VOLUME_roi20, VOLUME_roi29, VOLUME_roi22, VOLUME_roi04, VOLUME_roi40, VOLUME_roi37, VOLUME_roi19, VOLUME_roi16, VOLUME_roi17, VOLUME_roi38, VOLUME_roi36, VOLUME_roi13, VOLUME_roi03, VOLUME_roi12, VOLUME_roi11, VOLUME_roi34, VOLUME_roi06, VOLUME_roi21, VOLUME_roi08, VOLUME_roi24, VOLUME_roi10, VOLUME_roi14, VOLUME_roi30, VOLUME_roi32, VOLUME_roi07, VOLUME_roi28, VOLUME_roi23, VOLUME_roi31, VOLUME_roi27 |

**Table S3 |** Blood markers significantly associated with the brain-derived molecular AD dementia pseudotime.

**Markers** (all P<0.05, FDR-corrected; adjusted by age, sex and educational level)

Protein: CTF1 (Z=6.2); Metabolite: TG(51:5) (Z=-5.79); Protein: LRRN1 (Z=4.97); Metabolite: PC(32:4) (Z=4.67); Metabolite: PC-O(36:3) (Z=4.56); Metabolite: Gly (Z=-4.49); Metabolite: PC(37:7) (Z=4.19); Metabolite: PC(38:2) (Z=4.15); Protein: ITGA11|ITGB1 (Z=-4.13); Metabolite: TG(55:7) (Z=-4.12); Protein: DCAF12 (Z=4.12); Protein: ST8SIA2 (Z=4.08); Protein: ITGAV|ITGB3 (Z=-4.06); Protein: ACRV1 (Z=4.01); Protein: ERGIC1 (Z=3.96); Metabolite: PC(44:12) (Z=-3.96); Metabolite: AC(16:2-OH) (Z=-3.94); Protein: SEPTIN6 (Z=3.87); RNA: MCAT (Z=3.87); Protein: PLTP (Z=-3.86); Metabolite: SM(42:3) (Z=3.85); Protein: PIANP (Z=-3.81); Protein: ASCC2 (Z=-3.77); Protein: NPFF (Z=3.76); Metabolite: PC(40:8) (Z=-3.75); RNA: VAV3 (Z=3.75); Protein: OTULIN (Z=3.74); Protein: SWAP70 (Z=-3.73); Protein: DCAF11 (Z=3.72); Protein: NTM (Z=-3.71); Protein: C17orf78 (Z=3.7); Protein: PGAM2 (Z=-3.69); Protein: ATP6V1F (Z=-3.68); Metabolite: PC(39:7) (Z=-3.65); Protein: CSTB (Z=-3.65); Protein: OTUD3 (Z=3.63); Protein: CSMD2 (Z=-3.63); Protein: FCRL2 (Z=3.6); Protein: CCN5 (Z=3.6); RNA: FNDC3B (Z=3.58); Protein: LRRC3B (Z=-3.58); RNA: SVBP (Z=3.56); Metabolite: TG(56:6) (Z=-3.56); Protein: SDF2L1 (Z=-3.56); Protein: LTA4H (Z=3.55); Protein: CBR3 (Z=-3.55); Protein: SPSB1 (Z=3.55); Protein: TIE1 (Z=-3.54); Protein: ANTXR2 (Z=-3.54); Protein: KCNMB3 (Z=3.54); Protein: DNAJC1 (Z=-3.53); Protein: GSTA3 (Z=3.51); RNA: DNAJC27 (Z=3.51); Protein: SEMA6C (Z=3.49); RNA: NMRK1 (Z=3.49); Protein: MMEL1 (Z=3.48); Protein: CBLIF (Z=-3.48); Protein: EXOSC1 (Z=3.47); Protein: OGFR (Z=-3.47); Protein: NTS (Z=-3.47); Protein: IL22RA1 (Z=3.46); Protein: SNAI2 (Z=3.45); Protein: CFL2 (Z=-3.44); RNA: LPL (Z=3.44); Protein: CYBRD1 (Z=3.43); Protein: SLC35G2 (Z=3.43); Protein: STX12 (Z=-3.43); Metabolite: DG-O(34:1) (Z=3.41); Metabolite: CE(17:1) (Z=-3.41); Protein: MXRA8 (Z=-3.41); Protein: UMOD (Z=-3.4); Protein: CSF1R (Z=-3.38); Protein: FCAR (Z=3.37); Protein: TPPP3 (Z=-3.37); Protein: PAX8 (Z=3.36); RNA: ITGB1 (Z=3.35); Protein: SCARA3 (Z=3.35); Protein: SELENOW (Z=-3.35); Protein: PRKRA (Z=3.35); RNA: YTHDF3 (Z=3.33); Protein: RBP7 (Z=-3.33); Protein: NPTXR (Z=-3.33); Protein: AFAP1L2 (Z=-3.33); Protein: ANTXR1 (Z=-3.32); Protein: SEMA6B (Z=-3.32); Protein: BRD2 (Z=3.32); Protein: FAM24B (Z=3.32); Protein: ARL15 (Z=3.32); Protein: SIGLEC7 (Z=-3.31); Protein: CTNNA1 (Z=-3.31); Protein: CDC42EP4 (Z=-3.31); RNA: NUP93 (Z=3.3); Protein: HIF1AN (Z=3.3); RNA: SDHAP1 (Z=3.3); Protein: TPPP2 (Z=-3.29); Metabolite: DG(38:0) (Z=3.29); Protein: PGAM1 (Z=-3.29); Protein: SSBP1 (Z=-3.29); Metabolite: PC(40:7) (Z=-3.29); Protein: FN1 (Z=3.28); Protein: ACBD6 (Z=-3.28); Protein: PCSK1 (Z=-3.27); Protein: KLK8 (Z=-3.27); Protein: TFF2 (Z=-3.27); Protein: BEX4 (Z=3.27); Protein: PECR (Z=3.26); RNA: SIRPA (Z=3.26); Protein: TTC17 (Z=3.26); Protein: NPS (Z=-3.25); Protein: PIWIL1 (Z=3.25); RNA: GZF1 (Z=3.25); Protein: MSX2 (Z=3.24); Metabolite: PC(31:1) (Z=-3.24); RNA: PACSIN1 (Z=-3.24); Protein: ITIH2 (Z=3.24); Protein: TPSG1 (Z=3.23); RNA: RBM48 (Z=3.23); RNA: SULF1 (Z=3.23); Protein: RAB3D (Z=3.23); Protein: EGFR (Z=3.22); Metabolite: TG(50:4) (Z=3.22); Protein: PTK7 (Z=-3.2); Protein: SPC25 (Z=3.2); Protein: MLF1 (Z=3.2); Protein: MAMDC2 (Z=3.2); Protein: HS3ST5 (Z=3.19); Metabolite: CE(16:1) (Z=3.19); RNA: SDHAP1 (Z=3.19); Protein: METRNL (Z=-3.19); Protein: STK3 (Z=3.18); Protein: HENMT1 (Z=3.18); Protein: TRIB2 (Z=3.18); Protein: PIANP (Z=-3.18); RNA: ARID3B (Z=3.17); RNA: TLN1 (Z=3.17); RNA: RENBP (Z=3.17); RNA: CGNL1 (Z=-3.17); Protein: PTPRU (Z=-3.17); Protein: RAET1L (Z=-3.17); Protein: NRP1 (Z=-3.16); Protein: GEM (Z=3.16); Protein: GFAP (Z=3.16); Protein: TPPP (Z=3.16); RNA: PLA2G3 (Z=-3.16); Protein: SERPINH1 (Z=-3.15); Protein: EHMT2 (Z=-3.15); Protein: MGAT4C (Z=-3.14); RNA: KLF1 (Z=3.14); RNA: SLC25A45 (Z=-3.13); RNA: ING3 (Z=3.13); Protein: CLEC2D (Z=3.12); Protein: PLAU (Z=-3.12); Protein: DLG3 (Z=3.12); Protein: ABHD12 (Z=3.11); RNA: ZNF546 (Z=3.11); RNA: ZNF546 (Z=3.11); Protein: IGFBP4 (Z=-3.11); RNA: SLC19A1 (Z=3.11); Protein: UMODL1 (Z=3.1); Protein: PAGE4 (Z=3.1); Protein: CISD2 (Z=3.1); Protein: TREM1 (Z=-3.1); RNA: SRSF3 (Z=3.1); RNA: EPS8 (Z=3.1); RNA: SUPT16H (Z=3.09); Protein: TNC (Z=-3.09); Protein: HEXB (Z=-3.09); RNA: MEI1 (Z=3.09); Protein: KHSRP (Z=-3.08); Protein: CD209 (Z=-3.08); Protein: SELENBP1 (Z=3.08); Protein: LCE3B (Z=3.08); Metabolite: Asp (Z=-3.07); Protein: CENPW (Z=3.07); Protein: KYAT3 (Z=-3.07); RNA: (Z=3.07); Protein: CD5 (Z=-3.07); Protein: PSG6 (Z=3.07); Protein: PAPPA2 (Z=3.07); Protein: RS1 (Z=3.06); RNA: MX1 (Z=3.06); Protein: CANX (Z=3.06); Protein: BMP4 (Z=-3.05); Protein: BMP6 (Z=-3.05); Protein: SNRPF (Z=-3.04); Protein: ABI3 (Z=3.04); Protein: BAP18 (Z=-3.04); Protein: NPNT (Z=3.04); Protein: NPTXR (Z=-3.04); Protein: NFASC (Z=-3.04); Protein: GGPS1 (Z=3.04); RNA: SNX16 (Z=3.04); Protein: CAPNS2 (Z=-3.03); Protein: CD70 (Z=3.03); Metabolite: PC(35:4) (Z=-3.03); Protein: AMIGO2 (Z=-3.02); RNA: NCKAP5L (Z=3.02); RNA: PEX26 (Z=3.02); Protein: QPRT (Z=3.02); Protein: VPS29 (Z=-3.01); Protein: FCGR2B (Z=-3.01); RNA: CNGB3 (Z=-3.01); RNA: MAU2 (Z=3.01); Protein: SMPDL3A (Z=3.01); RNA: C10orf105 (Z=-3.01); RNA: LRRC37B (Z=-3.01); Protein: KLK8 (Z=-3); Protein: MYOC (Z=-3); Protein: CD300LF (Z=3); Protein: IL17A|IL17F (Z=3); Protein: IGFLR1 (Z=-3); Protein: IL22RA2 (Z=-3); RNA: C13orf46 (Z=2.99); RNA: C13orf46 (Z=2.99); Metabolite: PC-O(36:2) (Z=2.99); Protein: FATE1 (Z=2.99); Protein: ARHGAP45 (Z=-2.98); Protein: SEMA6D (Z=-2.98); RNA: LIAS (Z=2.98); Protein: NGFR (Z=-2.98); RNA: SEL1L (Z=2.98); Protein: PHGDH (Z=-2.97); Protein: ADGRE2 (Z=-2.97); Protein: THRA (Z=2.97); Protein: FAM20B (Z=-2.96); Metabolite: AC(4:0) (Z=-2.96); Protein: VSX1 (Z=2.96); Protein: None (Z=2.96); Protein: C17orf78 (Z=2.96); RNA: CERT1 (Z=2.96); Protein: MMP10 (Z=-2.95); Protein: GNAI3 (Z=-2.95); RNA: ILVBL (Z=2.95); Protein: ZPLD1 (Z=2.94); Protein: GAA (Z=-2.94); Protein: FN1 (Z=2.94); RNA: GULP1 (Z=2.94); Protein: POLR2J (Z=2.94); Protein: SYAP1 (Z=-2.94); RNA: LPAR2 (Z=-2.93); Protein: LYG1 (Z=-2.93); Protein: LEMD1 (Z=-2.93); Protein: YARS1 (Z=-2.93); RNA: PABPC4 (Z=2.93); RNA: CFAP410 (Z=2.93); RNA: SNAP47 (Z=2.92); Protein: FCGR3B (Z=-2.92); Protein: BAGE3 (Z=-2.92); RNA: STIL (Z=2.92); Protein: NFKBIA (Z=2.92); Protein: NNMT (Z=-2.92); Protein: BIRC7 (Z=2.91); Protein: XYLT2 (Z=2.91); Protein: ELMO1 (Z=-2.91); Protein: ADAM32 (Z=2.91); RNA: DNAJC16 (Z=2.91); Protein: NUBP2 (Z=2.91); Protein: KLF4 (Z=2.9); RNA: RPLP0 (Z=2.9); Protein: FANCF (Z=2.9); Protein: CYB5R2 (Z=-2.9); Protein: QSOX1 (Z=-2.89); Protein: SH3BGRL (Z=-2.89); Protein: APOA4 (Z=-2.89); Protein: CD14 (Z=-2.89); RNA: AKT2 (Z=2.89); Protein: NUDC (Z=-2.89); Protein: ARHGDIA (Z=2.89); Protein: CAB39L (Z=-2.89); Protein: VAPA (Z=-2.89); Protein: SLC5A8 (Z=2.88); Protein: GDAP1L1 (Z=2.88); RNA: ALOX12 (Z=2.88); Protein: ETNK2 (Z=2.88); Protein: LDHA (Z=-2.88); Protein: ABRAXAS1 (Z=2.88); Protein: METAP2 (Z=-2.88); Protein: NRDC (Z=2.87); RNA: RNASE4 (Z=2.87); Protein: SHC2 (Z=2.87); Protein: DCTPP1 (Z=-2.87); Protein: SEMA4F (Z=2.87); RNA: SLC5A6 (Z=2.87); RNA: IFIT3 (Z=-2.87); Protein: MMP10 (Z=-2.86); RNA: FAM53B (Z=2.86); Protein: CYREN (Z=-2.86); RNA: ORC3 (Z=2.86); Protein: RP9 (Z=-2.86); RNA: SAMD4B (Z=2.85); RNA: RASA4 (Z=2.85); Protein: SCAMP5 (Z=2.85); Protein: FTL (Z=2.85); RNA: PARP8 (Z=2.85); RNA: CDK16 (Z=2.85); Protein: CRELD1 (Z=-2.85); Protein: C1QTNF1 (Z=-2.85); Protein: MOCOS (Z=2.84); RNA: CCDC71L (Z=-2.84); RNA: GLB1L (Z=2.84); Metabolite: PC(32:6) (Z=2.84); RNA: CELF1 (Z=2.84); Protein: MRPS14 (Z=2.84); RNA: ABCA9 (Z=2.84); Protein: SORT1 (Z=2.84); Protein: FN1 (Z=2.84); Protein: RBP4 (Z=-2.84); RNA: AKAP10 (Z=2.84); Protein: STMN1 (Z=-2.83); RNA: KY (Z=2.83); Protein: ITPRIPL1 (Z=-2.83); Protein: UBE2M (Z=2.83); Protein: NEFL (Z=-2.83); Protein: CTSF (Z=2.83); Protein: SPRED1 (Z=2.82); Protein: ABHD14B (Z=2.82); Protein: PLOD2 (Z=-2.82); Protein: ATOX1 (Z=-2.82); Protein: ADM (Z=-2.82); Protein: PTGFRN (Z=-2.82); Metabolite: PC-O(40:7) (Z=2.81); Protein: DIAPH1 (Z=-2.81); Protein: PRKCG (Z=2.81); Protein: ALPG (Z=2.81); RNA: PRRC2C (Z=2.81); RNA: CASP10 (Z=-2.8); Protein: NLRP4 (Z=2.8); RNA: RNF150 (Z=2.8); Protein: ALKBH2 (Z=2.8); Protein: CASP10 (Z=-2.8); RNA: PIK3AP1 (Z=2.8); Protein: APOL3 (Z=-2.8); Protein: ETNK1 (Z=2.8); Protein: COA4 (Z=-2.8); RNA: PKD1P2 (Z=-2.8); Protein: LGMN (Z=-2.8); Protein: ZNF180 (Z=2.8); RNA: SLC25A22 (Z=2.79); Protein: CA11 (Z=-2.79); Protein: ACVRL1 (Z=-2.79); RNA: PPIAP82 (Z=-2.79); Protein: CD3G (Z=2.79); Protein: CCL24 (Z=2.79); Protein: FNDC4 (Z=2.79); Protein: SATB1 (Z=2.79); Protein: NTM (Z=-2.78); RNA: LARP4 (Z=2.78); RNA: SLC7A6 (Z=2.78); RNA: TLE1 (Z=2.78); Protein: KERA (Z=-2.78); RNA: RAB4A (Z=2.78); Protein: TMOD3 (Z=-2.78); Protein: POLE3 (Z=2.78); Protein: ADA2 (Z=2.77); Protein: SERF1A (Z=2.77); Protein: NPM2 (Z=2.77); Protein: DOK2 (Z=-2.77); RNA: STX5 (Z=2.77); RNA: ATF7 (Z=2.77); Protein: IL31RA (Z=2.77); Protein: UBA2 (Z=-2.77); Protein: TMPRSS6 (Z=2.77); Protein: PTPN2 (Z=2.77); Protein: SUDS3 (Z=2.77); RNA: LGALS3BP (Z=2.77); Protein: MYL5 (Z=2.77); RNA: YAF2 (Z=2.77); Protein: GCNT1 (Z=2.77); Protein: ADGRF1 (Z=2.76); RNA: EMCN (Z=2.76); Protein: SPINK4 (Z=-2.76); Protein: PTTG1 (Z=2.76); Protein: CT83 (Z=2.76); Protein: CERS5 (Z=2.76); RNA: NOC2LP2 (Z=-2.76); Protein: TNRC6B (Z=-2.76); RNA: ATPAF2 (Z=2.76); Protein: PCDH17 (Z=-2.75); Protein: ATP5PB (Z=-2.75); Protein: IL16 (Z=-2.75); RNA: DYNLL2 (Z=2.75); Protein: SMR3A (Z=2.75); Protein: SPINK9 (Z=-2.75); Protein: PLA2G2C (Z=2.75); RNA: TASOR2 (Z=2.75); RNA: ENTPD6 (Z=2.75); RNA: TRPM3 (Z=2.75); Protein: GGT5 (Z=-2.75); Protein: ADAM12 (Z=2.75); Protein: CSF1R (Z=-2.75); Protein: LUM (Z=-2.75); Protein: SCUBE3 (Z=-2.74); Protein: METAP1D (Z=2.74); Protein: C4A|C4B (Z=2.74); Protein: CDH17 (Z=-2.74); Protein: ADAMDEC1 (Z=2.74); RNA: YY1AP1 (Z=2.73); Protein: DEFB103A (Z=2.73); Protein: BMP4 (Z=2.73); RNA: ATL2 (Z=2.73); Protein: ISL1 (Z=2.73); Protein: FN1 (Z=2.73); RNA: ESYT2 (Z=2.73); Protein: SPON2 (Z=-2.73); Protein: C5orf38 (Z=2.73); RNA: KLHL5 (Z=2.73); Protein: MAP3K11 (Z=-2.72); Protein: CST5 (Z=-2.72); RNA: TIGD6 (Z=2.72); RNA: JAM2 (Z=2.72); Protein: SNX16 (Z=-2.72); Protein: FOXO1 (Z=-2.72); Protein: PLA2G2E (Z=2.72); Protein: UFC1 (Z=-2.71); Protein: FTH1|FTL (Z=2.71); RNA: IPCEF1 (Z=2.71); Protein: SERPINA3 (Z=2.71); Protein: LRFN3 (Z=2.71); RNA: ETHE1 (Z=2.71); Protein: FER (Z=-2.71); RNA: KIAA1109 (Z=2.71); RNA: HUWE1 (Z=2.71); Protein: SERPINA9 (Z=-2.7); Protein: PLG (Z=2.7); Protein: MB (Z=-2.7); RNA: GOLGA4 (Z=2.69); RNA: TCF12 (Z=2.69); Protein: CYB5R3 (Z=-2.69); Protein: INIP (Z=2.69); RNA: ABCB8 (Z=2.69); Protein: ACE2 (Z=2.69); Protein: ARL3 (Z=-2.69); RNA: HDAC7 (Z=2.69); RNA: FUT1 (Z=2.69); Protein: DPY30 (Z=-2.68); RNA: WDR45 (Z=2.68); Protein: CDK2|CCNA2 (Z=2.68); Protein: ZBTB16 (Z=2.68); Protein: SEMA3B (Z=2.68); Protein: PABPC4 (Z=-2.68); RNA: AMOTL1 (Z=2.68); Protein: TNXB (Z=-2.68); Protein: TMOD2 (Z=-2.68); Protein: None (Z=2.68); RNA: NAP1L4 (Z=2.67); RNA: NAP1L4 (Z=2.67); RNA: CNOT2 (Z=2.67); RNA: COG5 (Z=2.67); RNA: NFE2L1 (Z=2.67); Protein: RIPPLY1 (Z=2.66); RNA: TSPAN19 (Z=2.66); RNA: TTC21B (Z=2.66); Protein: GALNT16 (Z=-2.66); Protein: B3GAT3 (Z=-2.66); Protein: SAMSN1 (Z=-2.66); RNA: FNDC5 (Z=-2.66); Protein: NT5E (Z=-2.66); RNA: DNAL1 (Z=2.66); Metabolite: LPC-O(18:2) (Z=2.66); Protein: PTHLH (Z=-2.66); Protein: SYNGR3 (Z=2.66); Protein: AKT3 (Z=-2.65); Protein: IQCF3 (Z=2.65); RNA: SEC23B (Z=2.65); Protein: TNFRSF10A (Z=2.65); RNA: PDCD1 (Z=-2.65); Protein: GZMH (Z=2.65); RNA: WDR48 (Z=2.65); RNA: CDHR3 (Z=-2.65); RNA: DNAJB5 (Z=2.65); Protein: AKT1 (Z=-2.65); RNA: ZNF319 (Z=2.65); Protein: LRRC32 (Z=-2.65); RNA: CFAP298-TCP10L (Z=2.64); RNA: RNA5-8SP6 (Z=-2.64); Protein: DCLK3 (Z=2.64); Protein: SPINK7 (Z=-2.64); RNA: CDK15 (Z=2.64); Protein: ARL5B (Z=2.63); Protein: CLEC4G (Z=2.63); Protein: PTPRJ (Z=-2.63); Protein: FLRT1 (Z=2.63); Protein: DCP1B (Z=-2.63); RNA: KCTD7 (Z=2.63); Protein: PPCDC (Z=2.63); Protein: ADGRE5 (Z=-2.63); Protein: ADAM30 (Z=2.62); RNA: TRIM28 (Z=2.62); Protein: MPI (Z=-2.62); Protein: ATF1 (Z=2.62); Protein: SULT4A1 (Z=2.62); Protein: UBE2T (Z=2.62); Protein: MERTK (Z=-2.62); RNA: SYN1 (Z=-2.62); RNA: U2AF1 (Z=2.62); Protein: ADCYAP1 (Z=2.62); Protein: SERPING1 (Z=2.61); RNA: UIMC1 (Z=2.61); Protein: TACSTD2 (Z=2.61); Protein: BOC (Z=-2.61); Protein: RXFP1 (Z=2.61); RNA: PLOD3 (Z=2.61); RNA: STAG3L5P (Z=2.61); Protein: CT45A3 (Z=2.61); RNA: DCAF11 (Z=-2.6); RNA: DCAF11 (Z=-2.6); Protein: CTSE (Z=-2.6); Protein: PRDX5 (Z=2.6); RNA: DDX52 (Z=2.6); Protein: ING4 (Z=-2.6); RNA: CACNA1C (Z=2.6); RNA: (Z=2.6); RNA: MYO1G (Z=2.59); RNA: ITGAV (Z=2.59); Protein: PABPC3 (Z=-2.59); Protein: DEFB106A (Z=2.59); RNA: HIVEP1 (Z=-2.59); Protein: TMEM132D (Z=-2.59); Protein: RAB27A (Z=-2.59); RNA: MAEA (Z=2.59); Protein: NIF3L1 (Z=-2.59); Protein: ZKSCAN7 (Z=2.59); RNA: SMAD5 (Z=2.59); Protein: SEMA3G (Z=-2.59); Protein: VHL (Z=2.59); RNA: DNAJA4 (Z=2.58); Protein: SHC4 (Z=2.58); RNA: GGNBP2 (Z=2.58); RNA: GGNBP2 (Z=2.58); Protein: BLMH (Z=2.58); Protein: PMVK (Z=-2.58); Protein: GKN2 (Z=-2.58); RNA: SEMA3G (Z=-2.58); Protein: CD209 (Z=-2.58); Protein: USF2 (Z=2.58); RNA: ATF7IP2 (Z=-2.58); Protein: BICD1 (Z=2.58); RNA: ZNF524 (Z=2.57); RNA: FBXO38 (Z=2.57); Protein: ACAN (Z=2.57); RNA: RPL3 (Z=2.57); RNA: ZNF136 (Z=2.57); RNA: RABL6 (Z=-2.57); Protein: ELL2 (Z=-2.57); Protein: IGHG1 (Z=2.57); RNA: RNF19A (Z=-2.57); Protein: APBB1IP (Z=-2.57); Protein: TYMP (Z=-2.57); RNA: FMO6P (Z=-2.57); Protein: MRAP (Z=2.57); Protein: SEPTIN11 (Z=-2.57); Protein: PCSK2 (Z=2.56); RNA: MAX (Z=-2.56); Protein: GBP1 (Z=-2.56); Protein: LILRA4 (Z=-2.56); RNA: GSE1 (Z=2.56); RNA: AFF1 (Z=-2.56); RNA: ZNF670 (Z=2.56); RNA: PPP3CA (Z=2.56); RNA: KAT6B (Z=2.56); RNA: KAT6B (Z=2.56); RNA: BCLAF1 (Z=2.56); RNA: SELENOW (Z=-2.56); RNA: SNX13 (Z=2.56); RNA: SETD5 (Z=2.56); RNA: ANTXRLP1 (Z=2.56); Protein: NMU (Z=2.56); Protein: MASP1 (Z=2.56); RNA: SIGIRR (Z=2.55); Protein: None (Z=2.55); Protein: SH3PXD2B (Z=-2.55); RNA: RACK1 (Z=2.55); Protein: TRABD (Z=2.55); Protein: SND1 (Z=2.55); RNA: MTCO1P24 (Z=-2.55); RNA: DDX3X (Z=2.55); Protein: SDSL (Z=-2.55); Protein: SCPEP1 (Z=-2.55); Metabolite: LPC(17:0) (Z=-2.55); RNA: LRRC37A2 (Z=-2.55); Metabolite: PC(42:10) (Z=-2.55); RNA: FAM83H (Z=-2.55); Protein: FAM174A (Z=-2.55); Metabolite: PC(34:5) (Z=2.55); Protein: PRKAA1|PRKAB1|PRKAG1 (Z=2.55); Protein: PLAAT3 (Z=2.55); Protein: LEG1 (Z=2.54); Protein: PLA2G7 (Z=2.54); Protein: FBXL4 (Z=2.54); Protein: KLRF1 (Z=2.54); Protein: METAP1 (Z=-2.54); Protein: CCDC50 (Z=-2.54); RNA: MYO15B (Z=2.54); Protein: MAPKAPK3 (Z=-2.54); Protein: VASN (Z=-2.54); Protein: EPS8L2 (Z=-2.54); Protein: SETBP1 (Z=2.54); Protein: NUDT16 (Z=-2.54); Protein: IL12A|IL12B (Z=2.53); Metabolite: PC-O(42:4) (Z=2.53); Protein: PLAUR (Z=-2.53); Metabolite: CE(18:1) (Z=-2.53); Protein: LILRB3 (Z=2.53); Metabolite: PC-O(40:3) (Z=-2.53); RNA: NECAB1 (Z=2.53); Protein: APOH (Z=2.53); RNA: USP7 (Z=2.53); RNA: RRBP1 (Z=2.53); Protein: PGF (Z=2.53); Protein: FAS (Z=2.53); RNA: VAPB (Z=-2.52); Protein: SPAST (Z=2.52); RNA: ENO1 (Z=2.52); Protein: HSPA12A (Z=2.52); Protein: CST3 (Z=-2.52); Protein: PFN2 (Z=2.52); Metabolite: LPC-O(17:1) (Z=-2.52); Protein: BROX (Z=2.52); RNA: HSF1 (Z=2.52); RNA: HSF1 (Z=2.52); Protein: MRPL32 (Z=2.52); Protein: GLUD2 (Z=2.52); Protein: YME1L1 (Z=2.52); Protein: MAPKAPK5 (Z=2.51); RNA: SERF2 (Z=2.51); RNA: ITGA1 (Z=-2.51); Protein: EBP (Z=2.51); RNA: ATXN7L1 (Z=2.51); RNA: SNX13 (Z=2.51); Protein: FABP2 (Z=-2.51); RNA: N4BP2L2 (Z=2.51); Protein: RAB2B (Z=2.51); Protein: PLEKHO2 (Z=-2.51); Protein: SAT1 (Z=2.5); RNA: SLC41A1 (Z=-2.5); RNA: BCLAF1 (Z=2.5); RNA: COMMD2 (Z=2.5); Protein: GRN (Z=-2.5); RNA: IRF2 (Z=2.5); Protein: PAX4 (Z=2.5); Protein: TRIM27 (Z=2.5); RNA: TFRC (Z=2.5); RNA: ALG5 (Z=2.5); RNA: PIGO (Z=2.49); Protein: PCBP2 (Z=-2.49); RNA: CARMIL2 (Z=-2.49); Protein: KIR2DL4 (Z=2.49); Protein: DOCK2 (Z=2.49); Protein: FAHD2A (Z=-2.48); RNA: HSP90B1 (Z=2.48); RNA: LFNG (Z=2.48); RNA: INIP (Z=2.48); RNA: AGAP10P (Z=2.48); Protein: ITGA6 (Z=2.48); RNA: ZMYM2 (Z=2.48); RNA: FZR1 (Z=2.48); Protein: CRPPA (Z=2.48); RNA: ZFAND6 (Z=2.48); RNA: TAF2 (Z=2.48); Protein: HPGDS (Z=-2.48); RNA: MICOS10 (Z=2.47); Protein: PUS7 (Z=-2.47); RNA: TYK2 (Z=2.47); RNA: CARMIL3 (Z=-2.47); RNA: CARMIL3 (Z=-2.47); Protein: ZNF264 (Z=2.47); Protein: KDM4C (Z=-2.47); Protein: APEX2 (Z=2.47); RNA: GINS3 (Z=2.46); Protein: CCNB1IP1 (Z=2.46); RNA: TSPAN32 (Z=2.46); RNA: VPS29 (Z=2.46); RNA: TBC1D13 (Z=2.46); RNA: ABCF1 (Z=-2.45); RNA: ABCF1 (Z=-2.45); RNA: ABCF1 (Z=-2.45); Protein: FABP1 (Z=-2.45); RNA: ABHD16A (Z=2.45); RNA: NAALADL2 (Z=2.45); Protein: PIP (Z=2.45); RNA: RIN2 (Z=2.45); Protein: IL26 (Z=2.45); RNA: GPATCH8 (Z=2.45); RNA: PLAA (Z=2.45); RNA: CHST15 (Z=2.44); RNA: TM2D3 (Z=2.44); RNA: YTHDF3 (Z=2.44); Protein: ITGAL (Z=2.44); Protein: MCAM (Z=-2.44); Protein: F2 (Z=2.44); RNA: TEKT2 (Z=2.44); Protein: OBP2A (Z=2.44); RNA: SNRNP40 (Z=2.44); RNA: LMLN2 (Z=2.43); RNA: CROT (Z=-2.43); RNA: SAMD9 (Z=-2.43); RNA: DIPK2A (Z=-2.43); Protein: FAM171A2 (Z=2.43); RNA: TST (Z=2.43); Protein: MICOS10 (Z=2.43); Protein: IL1RAPL2 (Z=2.43); RNA: FAM135A (Z=2.42); RNA: HMGB3P9 (Z=2.42); Protein: PLCD1 (Z=-2.42); RNA: TRIM14 (Z=2.42); Metabolite: PC(43:6) (Z=2.42); RNA: ZNF318 (Z=2.41); RNA: NFATC1 (Z=2.41); RNA: CDK5RAP2 (Z=2.4); RNA: TUBB (Z=2.4); RNA: JUP (Z=2.4); Protein: PEX26 (Z=2.4); Protein: C1QBP (Z=2.39); RNA: PEX26 (Z=2.39); Protein: PRSS57 (Z=-2.38); RNA: GNAI2 (Z=2.38); RNA: C8orf33 (Z=2.38); Protein: ELK1 (Z=-2.37); RNA: CHD2 (Z=2.36); Protein: ZNF566 (Z=2.35); RNA: SBDSP1 (Z=2.34); Protein: LTB4R (Z=-2.34)

*Z values are relative correlations (i.e., # std over randomized values).

**Table S4.** Post-hoc subtype-subtype comparisons on neuroticism, conscientiousness, depressive symptoms, loneliness, and purpose in life.

| **Measure** | **Subtype-subtype comparisons** | **Difference in means** | **95% CI** | | ***P*** |
| --- | --- | --- | --- | --- | --- |
| **Depressive symptoms** | **0 – 1** | **0.34 (0.01)** | **-0.63** | **-0.06** | **0.009** |
|  | 1 – 2 | 0.14 (0.01) | -0.17 | 0.45 | 1.00 |
|  | 1 – 3 | 0.09 (0.01) | -0.41 | 0.23 | 1.00 |
|  | 0 – 2 | 0.20 (0.01) | -0.50 | 0.09 | 0.394 |
|  | 2 – 3 | 0.23 (0.01) | -0.56 | 0.09 | 0.34 |
|  | **0 – 3** | **0.44 (0.01)** | **-0.73** | **-0.14** | **<0.001** |
| **Neuroticism** | **0 – 1** | **2.38 (0.05)** | **-3.74** | **1.02** | **<0.001** |
|  | 1 – 2 | 0.70 (0.06) | -0.79 | 2.19 | 0.139 |
|  | 1 – 3 | 1.16 (0.06) | -0.37 | 2.69 | 0.270 |
|  | **0 – 2** | **1.68 (0.05)** | **-3.09** | **-0.27** | **0.010** |
|  | 2 – 3 | 0.46 (0.05) | -1.10 | 2.02 | 1.000 |
|  | 0 – 3 | 1.22 (0.05) | -2.64 | 0.20 | 0.139 |
| **Loneliness** | **0 – 1** | **0.21 (0.07)** | **-0.39** | **-0.02** | **0.022** |
|  | 1 – 2 | 0.03 (0.08) | -0.23 | 0.18 | 1.00 |
|  | 1 – 3 | 0.13 (0.08) | -0.08 | 0.33 | 0.64 |
|  | **0 – 2** | **0.23 (0.07)** | **-0.42** | **-0.04** | **0.007** |
|  | 2 – 3 | 0.15 (0.07) | -0.05 | -0.36 | 0.31 |
|  | 0 – 3 | 0.08 (0.07) | -0.26 | 0.11 | 1.00 |
| **Conscientiousness** | 0 – 1 | 0.78 (0.05) | -0.56 | 2.12 | 0.75 |
|  | 1 – 2 | 0.89 (0.06) | -0.62 | 2.40 | 0.72 |
|  | **1 – 3** | **1.58 (0.06)** | **0.02** | **3.14** | **0.046** |
|  | **0 – 2** | **1.67 (0.05)** | **0.23** | **3.11** | **0.013** |
|  | 2 – 3 | 0.69 (0.05) | -0.96 | 2.33 | 1.00 |
|  | **0 – 3** | **2.36 (0.05)** | **0.89** | **3.82** | **<0.001** |
| **Purpose in life** | 0 – 1 | 0.11 (0.05) | -0.02 | 0.24 | 0.18 |
|  | 1 – 2 | 0.08 (0.06) | -0.07 | 0.23 | 0.90 |
|  | 1 – 3 | 0.04 (0.06) | -0.11 | 0.19 | 1.00 |
|  | **0 – 2** | **0.19 (0.05)** | **0.06** | **0.33** | **0.001** |
|  | 2 – 3 | 0.04 (0.06) | -0.19 | 0.10 | 1.00 |
|  | **0 – 3** | **0.15 (0.05)** | **0.01** | **0.28** | **0.021** |

***** We used Bonferroni’s test for mean score.
